# THE INSECTICIDAL POTENTIAL OF *BACILLUS CEREUS* GROUP STRAINS FROM INSECT-DENSE REGIONS OF THE UNITED KINGDOM

**DOI:** 10.64898/2026.09.11.750885

**Authors:** Josefin Blom, Vishnu Raghuram, Ben Raymond, Laura M. Carroll

## Abstract

Entomopathogenic *Bacillus cereus* group members (e.g. *B. thuringiensis*) are arguably the most important source of biopesticides globally and are widely used for industrial-scale crop protection. However, the application of biopesticidal *B. cereus* group strains is currently threatened by increasing resistance among insects, necessitating the development of novel insecticides. Here, we used whole-genome sequencing to characterize 13 novel *B. cereus* group strains from sites with high-density insect populations, including six isolates from an insect cadaver. Overall, diverse *B. cereus* group members were observed, encompassing three *panC* groups (II, IV and VI) and seven sequence types. Five strains possessed insecticidal Cry toxin-encoding genes with predicted target ranges that included insect species present at the respective sampling locations. Moreover, the genomes were computationally mined for biosynthetic gene clusters (BGCs; clusters of genes governing secondary metabolite synthesis). In total, 274 BGCs were observed, of which the majority (170 BGCs; 62.04%) did not cluster with known BGCs. Taken together, our study indicates that sampling *B. cereus* group bacteria from insect-rich ecological niches may yield Cry toxins against the self-same insects, as well as novel and unexplored BGCs, suggesting that targeted sampling efforts could generate compound leads for biopesticide development.

## INTRODUCTION

The *Bacillus cereus* group (*B. cereus sensu lato*) is a species complex of Gram-positive, spore-forming bacteria that are nearly ubiquitous in nature (Stenfors Arnesen, Fagerlund, and Granum 2008, Mondol, Shin, and Islam 2013, Ehling-Schulz, Frenzel, and Gohar 2015). In addition to their wide prevalence in soil and marine environments, *B. cereus* group members also occupy niches in vertebrate and invertebrate hosts, such as animals and insects (Ehling-Schulz, Frenzel, and Gohar 2015). A notable example of an invertebrate-dwelling *B. cereus* group member is *B. thuringiensis*, which due to its insecticidal properties is widely used as a biopesticide (here, *Bt* will refer to commercial *Bt* strains used as crop protection agents, while the broader ensemble of entomopathogenic *B. cereus* group strains will be referred to as insecticidal *B. cereus* group strains; Ehling-Schulz, Frenzel, and Gohar 2015, Ragasruthi *et al*. 2024). In 2011, *Bt* alone represented approximately 2% of the insecticidal market, and six years later, there were >98 *Bt*-based spray formulations marketed for crop protection in the United States (Bravo *et al*. 2011, Jouzani, Valijanian, and Sharafi 2017). Additionally, *Bt*-derived genes have been used in transgenic versions of rice, corn, cotton and soybean crops, with the success of the transgenic crops surpassing that of the spray formulations (Bravo *et al*. 2011, Jouzani, Valijanian, and Sharafi 2017). Taken together, *Bt* is currently the most important biopesticide worldwide and considered safer than its chemical counterparts. However, its future is precarious given the increase in resistance among insects to *Bt*-based formulations, for which there are no proposed solutions (Maagd de, Bravo, and Crickmore 2001, Bode 2009, Bravo *et al*. 2011, 2013, Jouzani, Valijanian, and Sharafi 2017).

Biopesticidal *B. thuringiensis* derives its entomopathogenic activity from crystal proteins produced during the sporulation phase (Bravo *et al*. 2011, 2013, Jouzani, Valijanian, and Sharafi 2017). These crystal proteins, also referred to as δ-endotoxins, induce death via the perforation of insect larval midguts upon ingestion of the bacterial spores. In addition to the two families of δ-endotoxins (Cry and Cyt), the bacteria may also express toxins (Vips) during the vegetative phase. The combination of Cry, Cyt and Vip proteins (henceforth referred to as “*Bt*-associated toxins”) determine potential insect targets, with commercial *Bt* formulations providing protection against Lepidoptera, Coleoptera, Diptera and nematodes. The modification of known and the discovery of new δ-endotoxins could lead to the development of alternative *Bt*-based formulations (Bravo *et al*. 2011, 2013, Jouzani, Valijanian, and Sharafi 2017).

While the *Bt*-associated toxins are relatively well-known and researched, other virulence factors produced by entomopathogenic bacteria are less studied (la Fuente-Salcido de, Casados-Vázquez, and Barboza-Corona 2013). This includes secondary metabolites, which can aid adaptation to specific ecological niches and confer diverse bioactivities, e.g. antibacterial and insecticidal activity (Tyc *et al*. 2017, Mullowney *et al*. 2023). Examples include thuringiensin, an entomopathogenic adenine nucleoside oligosaccharide that acts as an RNA polymerase inhibitor to suppress insect metabolism (Levinson *et al*. 1990, Liu XY *et al*. 2010), and zwittermicin A, a linear aminopolyol that can amplify the insecticidal activity of *Bt*-strains (Kevany, Rasko, and Thomas 2009). Prior studies investigating the genetic determinants of secondary metabolite synthesis (biosynthetic gene clusters; BGCs) among the *B. cereus* group have highlighted the underused biosynthetic potential of the species complex (Zhao and Kuipers 2016, Grubbs *et al*. 2017, Xia *et al*. 2022, Vater *et al*. 2023, Yin *et al*. 2023, Blom *et al*. 2025), indicating that undiscovered secondary metabolites produced by insecticidal *B. cereus* group strains could exhibit entomopathogenic activity and be of value for biocontrol applications.

High insect density has been linked to an increased prevalence of insecticidal members of the *B. cereus* group, and consequently insect-dense areas and insect cadavers have historically proved to be valuable sources of biopesticidal strains (Dulmage 1970, Barjac de 1978, Martínez *et al*. 2004, Raymond *et al*. 2010). Here, we employ whole-genome sequencing to study 13 novel *B. cereus* group strains isolated from three insect-rich sites in the United Kingdom (UK) with no prior *Bt* application. By harnessing *in silico* methods for the detection of *Bt*-associated toxins and BGCs, we aim to provide a comprehensive overview of the virulence factors governing interactions between the *B. cereus* group strains, potential insect hosts and their environment. Finally, to gain insight into the epidemiology of these strains, we compare our genomes to high-quality, publicly available assembled *B. cereus* group genomes (*n* = 2890). Overall, we hope to provide a first glimpse into novel compound leads, which could be used to combat the growing resistance among pests to existing *Bt*-based formulations and transgenic crops.

## MATERIALS AND METHODS

### Isolation of *B. cereus* group strains

Isolates were recovered from soil, insect or leaf material from three localities in the UK with high-density, herbivorous insect populations. All bacteria were isolated on *B. cereus-* specific agar (Oxoid,UK) using previously described methods (Raymond *et al*. 2010); in brief, leaf samples were vortexed in sterile saline and sharp sand before plating the leaf wash at 30°C. All soil samples were diluted 10- and 100-fold in saline and pasteurized at 60°C before plating out.

The site at Silwood Park (Bracknell Forest) was part of a study of the dynamics of *B. cereus* group bacteria on broad-leaved dock (*Rumex obstusifolius*) heavily colonized by the green dock leaf beetle *Gastrophysa viridula* (De Geer; Chrysomelidae; Collier *et al*. 2005). The Shinfield site is part of the University of Reading’s agricultural research station and isolates were collected during a study of insect pests on *Brassica* crops (Staley *et al*. 2010). All samples were derived from a single Lepidopteran cadaver, likely to be a *Pieris* spp larva. The Orkney sites are lowland coastal heath communities dominated by *Calluna vulgaris* and were characterized by frequent outbreak densities of Geometrid caterpillars, predominantly winter moth *Operophtera brumata* (Graham *et al*. 2004). The isolates were derived from the soil/leaf litter layer in three localities listed by Graham *et al*. (2004): Hundland (HU), Linnadale (LI) and Settiscarth (SET).

### Phenotypic characterization

The isolates from Bracknell Forest and the Orkney Islands (except for 243879_SET22) were stained for *Bt*-associated toxins using a previously described protocol (Raymond *et al*. 2010).

### Whole-genome sequencing (WGS), data pre-processing, *de novo* assembly, annotation and quality control

All isolates, provided as live bacterial samples by BR, were sequenced by Microbes NG (https://microbesng.com/whole-genome-sequencing/) in 2023. Briefly, libraries were prepared using the Nextera XT Library Prep Kit (Illumina, USA) and sequenced on a NovaSeq6000 (Illumina, 2 x 250 bp paired-end reads).

The raw sequencing reads for all strains (*n* = 13) were run through the Bactopia pipeline (v3.0.1; Supplementary Table S1; Petit III and Read 2020) for WGS pre-processing, *de novo* assembly, annotation and quality control. Briefly, the raw Illumina reads were quality controlled using FastQC (v0.12.1; https://www.bioinformatics.babraham.ac.uk/projects/fastqc/), fastp (v0.23.4; Chen *et al*. 2018) and lighter (v1.1.2; Song, Florea, and Langmead 2014). The reads were thereafter assembled with Skesa wrapped in Shovill (v1.1.0; https://github.com/tseemann/shovill; Souvorov, Agarwala, and Lipman 2018). Subsequently, the *de novo* assembled genomes were annotated using Prokka (v1.14.6; Seemann 2014) and AMRFinderPlus (v3.12.8, database v2024-01-31.1; Feldgarden *et al*. 2019), and assigned to sequence types (STs) using mlst (v2.23.0; https://github.com/tseemann/mlst; Jolley and Maiden 2010). Assemblies were quality controlled with CheckM (v1.2.2; Parks *et al*. 2015) and QUAST (v5.2.0; Gurevich *et al*. 2013) using the Bactopia Tools modules with default settings (Supplementary Table S2; Petit III and Read 2020). All assembled genomes had CheckM completeness of >99% and contamination <1%, and were therefore deemed to be of high quality. For additional quality control, MultiQC (v1.16; Ewels *et al*. 2016) was run on the output of FastQC, produced by Bactopia (Supplementary Table S3).

### Taxonomic classification, sequence typing and virulence factor detection

For each of the 13 *de novo* assembled genomes generated in this study (referred to hereafter as “novel strains”), species were assigned according to the Genome Taxonomy Database (GTDB) classification via the pre-defined subworkflow for GTDB-Tk (v2.3.2; Chaumeil *et al*. 2020), present in Bactopia Tools (Petit III and Read 2020), with GTDB release R214 specified (accessed the 20^th^ of March 2024).

Using the BTyper3 (v3.4.0; Carroll, Cheng, and Kovac 2020, Carroll, Wiedmann, and Kovac 2020) subworkflow from Bactopia Tools, the novel strains were also assigned to (i) sequence types (STs) within PubMLST’s seven-gene multi-locus sequence typing scheme for “*B. cereus*” (Jolley and Maiden 2010), (ii) pantoate-beta-alanine ligase gene (*panC*) groups (I-VIII) and (iii) taxonomic units within the 2020 *B. cereus* group Genomospecies-Subspecies-Biovar (GSB) taxonomy (Carroll, Wiedmann, and Kovac 2020). Additionally, BTyper3 was used for the detection of virulence factors, including *Bt*-associated toxin genes (Supplementary Table S4). BTyper3 (v3.4.0; not part of Bactopia Tools) was similarly used to detect virulence factors in all publicly available genomes included in the maximum-likelihood phylogenies (*n* = 2461 genomes, including the outgroups; described below; Supplementary Table S5).

A previously published table, mapping *Bt*-associated toxin gene families to predicted insect targets, was downloaded to assess potential insecticidal activity among the strains based on the detected *Bt*-associated toxin genes (Table 2 from Jouzani, Valijanian, and Sharafi 2017). More insect target specificity files for Cry toxins were downloaded from the Bacterial Pesticidal Protein Database (BPPRC; accessed on the 9^th^ of January 2026; Crickmore *et al*. 2021). Briefly, Cry genes detected by BTyper3 were entered as search terms in BPPRC. If the BTyper3 results exhibited some uncertainty (e.g. Cry1Ac1/Cry1Ac10/Cry1Ac7/Cry1Ac8/Cry1Ac9), the more general Cry-family was used as a search term (e.g. Cry1Ac). The general Cry-family was also entered if the initial search failed to yield results, to cast a broader net. Any BTyper3 result of the format XXX-like (e.g. Cry1-like) was ignored. The resulting insect target specificity tables were downloaded as Excel files and examined for mentions of the relevant insect species.

### Average nucleotide identity (ANI)-based genome analysis

FastANI (v1.33; Jain *et al*. 2018) was used to compute ANI values between all 13 novel strains in an all-versus-all approach (*n* = 169 comparisons; Supplementary Figure S1; Supplementary Table S6). Subsequently, the pairwise ANI values were used to generate an ANI-based dendrogram with bactaxR (v0.2.3; Jain *et al*. 2018, Carroll, Wiedmann, and Kovac 2020). ANI values were also computed between the novel strains and publicly available genomes (described below) within the same 2020 GSB species.

### BGC mining and gene cluster family delineation

BGCs were detected in the 13 novel strains using: (i) antiSMASH (v8.0.4; Blin *et al*. 2025), a stringent, rule-based BGC detection tool, and (ii) GECCO (v0.9.8; Carroll *et al*. 2021), a machine learning-based tool which prioritizes novel BGCs. For (i) antiSMASH, the resulting region GenBank files were broken into individual protoclusters (*n* = 196 protoclusters), as the protocluster unit represents BGCs with a single product type (Blin *et al*. 2019). Thereafter, the detected BGCs (*n* = 78 for GECCO; *n* = 196 for antiSMASH) were concatenated with the Minimum Information about a Biosynthetic Gene cluster (MIBiG) database (v4.0; *n* = 2636 BGCs; Zdouc *et al*. 2024), to enable comparisons with previously known, validated BGCs. The resulting dataset (*n* = 2910 BGCs) was subsequently clustered into gene cluster families (GCFs) using IGUA (Larralde *et al*. 2025; Supplementary Table S7).

The biosynthetic classes of the BGCs derived from the novel strains were extracted from the GenBank records produced by antiSMASH (v8.0.4) and GECCO (v0.9.8). To ensure conservative biosynthetic class labels, classes associated with higher levels of uncertainty (i.e. “Unknown” and “Others”), as well as hybrids beyond the well-established non-ribosomally synthesized peptide-polyketides (NRP-PKS) class, were merged and labelled as “Others”.

To investigate the presence and completeness of the zwittermicin A BGC among public, biopesticidal strains, the commercial ST8 biopesticide strains from the publicly available genomes (described below; *n* = 12 strains) were also mined for BGCs. Briefly, antiSMASH and GECCO were used as above, with the exception that antiSMASH region files were not split into protoclusters.

Comparative gene cluster plots were generated using LoVis4u (v0.1.6; Egorov and Atkinson 2025) and clinker (v0.0.12; Gilchrist and Chooi 2021).

### Construction of maximum-likelihood phylogenies

To compare the 13 novel strains to publicly available genomes, a previously published dataset of genomes (*n* = 2890) and associated metadata were downloaded (accessed 26^th^ of August 2024; Carroll *et al*. 2022). The novel and the publicly available genomes were grouped based on their 2020 GSB species assignment (Carroll, Wiedmann, and Kovac 2020), thereby forming three datasets: *B. cereus s.s.* genomes (*n* = 1381), *B. mycoides* genomes (*n* = 195) and *B. mosaicus* genomes (*n* = 895).

Maximum-likelihood phylogenies were constructed for each dataset (*n* = 3), using the same workflow. Briefly, a publicly available genome was included as an outgroup (Supplementary Table S8). To identify core genes and produce core gene alignments for each dataset and its outgroup, Panaroo (v1.3.4; Tonkin-Hill *et al*. 2020) was used. For rapid extraction of single nucleotide polymorphisms (SNPs), the resulting core gene alignments were supplied as input for SNP-sites (v2.5.1; Page *et al*. 2016). IQ-TREE (v.2.2.5; Tavarc 1986, Yang 1995, Thi Hoang *et al*. 2017, Minh *et al*. 2020) was thereafter used to compute maximum-likelihood phylogenies based on the core SNPs. The generated phylogenies were rooted using the outgroup genomes, before the outgroups were omitted for readability. Visualization and annotation of the phylogenies were done using ggtree (v3.10.1; Yu *et al*. 2017), ape (v5.7-1; Paradis and Schliep 2019), phytools (v2.3-0; Revell 2024), ggnewscale (v0.4.10; https://github.com/eliocamp/ggnewscale) and phangorn (v2.11.1; Schliep 2011).

### Construction of ST8 maximum-likelihood phylogeny

Given that five out of the 13 novel strains were assigned to ST8, a widely known lineage containing several strains exploited in biopesticides, we decided to explore ST8 in closer detail. A separate phylogeny was created for ST8, including both novel ST8 genomes (*n* = 5) and publicly available ST8 genomes (*n* = 117). Firstly, ANI values were computed for all ST8 genomes (n = 122 genomes), using an all-versus-all approach (described above). Based on the ANI analysis, there was an outlier among the ST8 genomes (Supplementary Figure S2); this outlier genome was excluded, resulting in 121 ST8 genomes used in further analysis. Thereafter, snippy (v4.6.0; https://github.com/tseemann/snippy) was used to create a core SNP alignment based on assembled genomes, with the medoid genome (FBO_2016_Bt_food_EU_France_16SBCL380_SAMN12147278_SRR9600166_GTDBbombys epticus_GSBcereus_ST8_GroupIV_Bonis2021; from the ANI-based analysis using bactaxR with a threshold of 99.5 ANI) used as a reference genome. SNP-sites (v2.5.1; Page *et al*. 2016) was used to extract the constant sites, after which the phylogeny was created using IQ-TREE (v2.2.5; Kalyaanamoorthy *et al*. 2017, Minh *et al*. 2020). An attempt was made to detect recombination using Gubbins (v3.4; Croucher *et al*. 2015), but Gubbins was unable to run successfully, likely due to the alignment being comprised of many highly similar genomes. As such, the clean, full alignment file generated by snippy was supplied to snp-dists (v0.8.2; https://github.com/tseemann/snp-dists) to calculate core SNP distances (Supplementary Table S9).

To investigate how closely related our novel ST8 strains were to public biopesticidal and environmental ST8 strains, a SNP haplotype network was constructed for a ST8 subset. This subset consisted of biopesticidal and environmental strains from the publicly available dataset (*n* = 28 genomes), our five novel strains, and two classical, biopesticidal ST8 strains for reference (HD-1, NCBI RefSeq Assembly GCF_000710255.1; HD-73, NCBI RefSeq Assembly GCF_000338755.1; Liu G *et al*. 2013, Day *et al*. 2014). Initially, a core SNP alignment was constructed for the ST8 subset (*n* = 35 genomes) as for the full ST8 dataset, with the following alteration: the reference genome was 243884_A2m47, as ANI-based analysis determined this to be the medoid genome for the ST8 subset. The core SNP alignment was used as input to create a SNP haplotype network in R, using the packages ape (v5.7-1; Paradis and Schliep 2019) and pegas (v1.3; Paradis 2010).

### Statistical analysis, data visualization and reproducibility

Statistical analysis and plot construction were conducted in R (v4.3.1) using the following packages: phytools (v2.3-0; Revell 2024), tidyverse (v2.0.0; Wickham *et al*. 2019), ape (v5.7-1; Paradis and Schliep 2019), ggtree (v3.10.1; Yu *et al*. 2017), ggsankey (v0.0.99999; https://github.com/davidsjoberg/ggsankey), maps (v3.4.2; https://github.com/adeckmyn/maps), ComplexHeatmap (v2.18.0; Gu, Eils, and Schlesner 2016), mapdata (v2.3.1; https://github.com/cran/mapdata), circlize (v0.4.16; Gu *et al*. 2014), dendextend (v1.17.1; Galili 2015), pegas (v1.3; Paradis 2010) and bactaxR (v0.2.3; Carroll, Wiedmann, and Kovac 2020). Principal component analysis plots were created using besPLOT (v1.0.2; Carroll, Huisman, and Wiedmann 2020). Plots were exported as .svg files using the svglite (v2.1.3; https://github.com/r-lib/svglite) and ggpubr (v0.6.0; https://github.com/kassambara/ggpubr) packages.

The Bactopia pipeline, including subworkflows and modules, and the BGC mining relied on Nextflow (v24.04.2; Tommaso Di *et al*. 2017) for reproducibility.

### Data availability

NCBI accession numbers for the novel strain genomes generated in this study are available in Table 1. Raw data, scripts and all code (including full command lines) associated with the publication are available at Zenodo DOI: https://doi.org/10.5281/zenodo.22045136.

**Table 1.** Metadata associated with the novel strains.

| Novel strain | NCBI BioSample Accession | Location | Isolation year | GTDB species | Bt genes | Prevalent insect species | Additional metadata |
| --- | --- | --- | --- | --- | --- | --- | --- |
| 243877_HUI2 | SAMN62739943 | Orkney Islands | 2013 | *Bacillus_A bombysepticus* | Yes | Winter moth (*O. brumata* (L.)) | Isolated from soil near Lepidoptera outbreaks |
| 243878_LI52 | SAMN62739944 |  |  | *Bacillus_A bombysepticus* | Yes |  |  |
| 243879_SET22 | SAMN62739945 |  |  | *Bacillus_A mycoides* | No |  |  |
| 243881_A1g9 | SAMN62739946 | Bracknell Forest | 2002 | *Bacillus_A cereus_U* | No | Green dock beetle (*G. viridula*) | Isolated from grass vegetation in proximity to *R. obtusifolius* foliage |
| 243882_A2m21 | SAMN62739947 |  |  | *Bacillus_A bombysepticus* | Yes |  | Isolated from mature leaves of *R. obtusifolius* foliage |
| 243883_A2m19 | SAMN62739948 |  |  | *Bacillus_A bombysepticus* | Yes |  |  |
| 243884_A2m47 | SAMN62739949 |  |  | *Bacillus_A bombysepticus* | Yes |  |  |
| 243885_shin1 | SAMN62739950 | Reading | 2007 | *Bacillus_A cereus* | No | Lepidoptera | Isolated from a single caterpillar cadaver at an experimental farm |
| 243886_shin2 | SAMN62739951 |  |  | *Bacillus_A mycoides* | No |  |  |
| 243887_shin3 | SAMN62739952 |  |  | *Bacillus_A wiedmannii* | No |  |  |
| 243888_shin4 | SAMN62739953 |  |  | *Bacillus_A mycoides* | No |  |  |
| 243889_shin5 | SAMN62739954 |  |  | *Bacillus_A mycoides* | No |  |  |
| 243890_shin6 | SAMN62739955 |  |  | *Bacillus_A mycoides* | No |  |  |

### AI disclosure

ChatGPT (GPT-5.5, OpenAI) was used to assist the writing of the Python script for splitting antiSMASH region GenBank files into protocluster GenBank files (https://zenodo.org/records/22045136/files/antismash_converter.py?download=1Cpreview=1).

## RESULTS

### Diverse *B. cereus* group strains were observed at UK locations with high-density insect populations

*B. cereus* group strains (*n* = 13) were isolated from three sites within the UK between the years of 2002 and 2013 (Table 1; Figure 1). The isolated strains underwent WGS and genome assembly to produce 13 high-quality, *de novo* assembled genomes.

**Figure 1.**
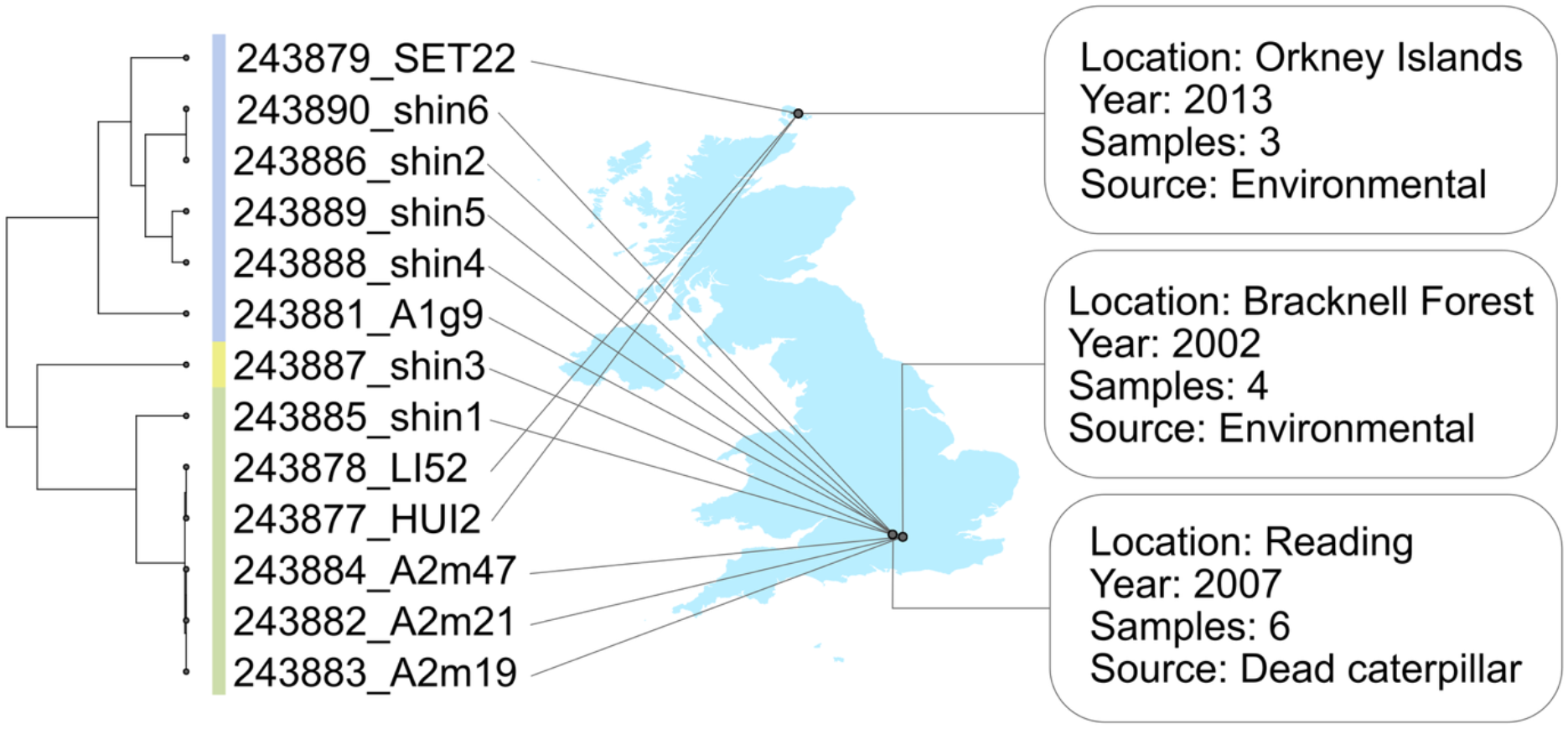
B. cereus group strains (n = 13) from UK sites with high-density insect populations are diverse. The left-hand dendrogram is based on average nucleotide identity (ANI), with branch lengths representing ANI distances. For readability, branch length ANI-values can be observed in the supplementary material (Supplementary Figure S1). The colour strip to the right of the dendrogram denotes panC Group assignment per BTyper3: blue = panC Group VI; yellow = panC Group II; green = panC Group IV.Alt. text: Graphical representation of the diversity of the novel B. cereus group strains, mapped onto the United Kingdom and with associated metadata.

To minimize any of the taxonomic ambiguities associated with species assignments within the *B. cereus* group (Carroll, Wiedmann, and Kovac 2020), all strains were characterized using (i) BTyper3’s *panC* classification, (ii) the Genome Taxonomy Database (GTDB) taxonomy, (iii) BTyper3’s 2020 Genomospecies-Subspecies-Biovar (GSB) taxonomy and (iv) the PubMLST seven-gene MLST scheme for “*B. cereus*” (Figure 2). For (i) *panC* group assignment, the eight-group classification employed by BTyper3 (Carroll, Cheng, and Kovac 2020, Carroll, Wiedmann, and Kovac 2020) was used. Overall, *panC* Groups IV and VI were the most prominent (*n* = 6 strains each), while only one genome was assigned to *panC* Group II (Figure 2; Supplementary Table S4). Based on (ii) the GTDB taxonomy, the 13 strains encompassed five GTDB species. The *panC* Group IV and VI genomes both spanned two GTDB species each, while the *panC* Group II genome was assigned to GTDB species *Bacillus_A wiedmannii*. According to (iii) the 2020 GSB taxonomy, the strains encompassed three species, with one *panC* group represented per GSB species. Only five genomes were positive for *Bt*-associated toxin genes, as determined via *in silico* methods, all belonging to *panC* Group IV. These genomes would thus be denoted as *B. cereus s.s.* biovar Thuringiensis per the 2020 GSB taxonomy (Carroll, Wiedmann, and Kovac 2020). None of the strains were predicted to possess genes encoding anthrax toxins or cereulide synthetase (i.e. genes responsible for the production of the emetic toxin cereulide; Supplementary Table S4). For (iv) the PubMLST MLST scheme, seven STs were represented among 12 out of the 13 strains; the last genome could not be assigned to an ST (Figure 2; Supplementary Table S4).

**Figure 2.**
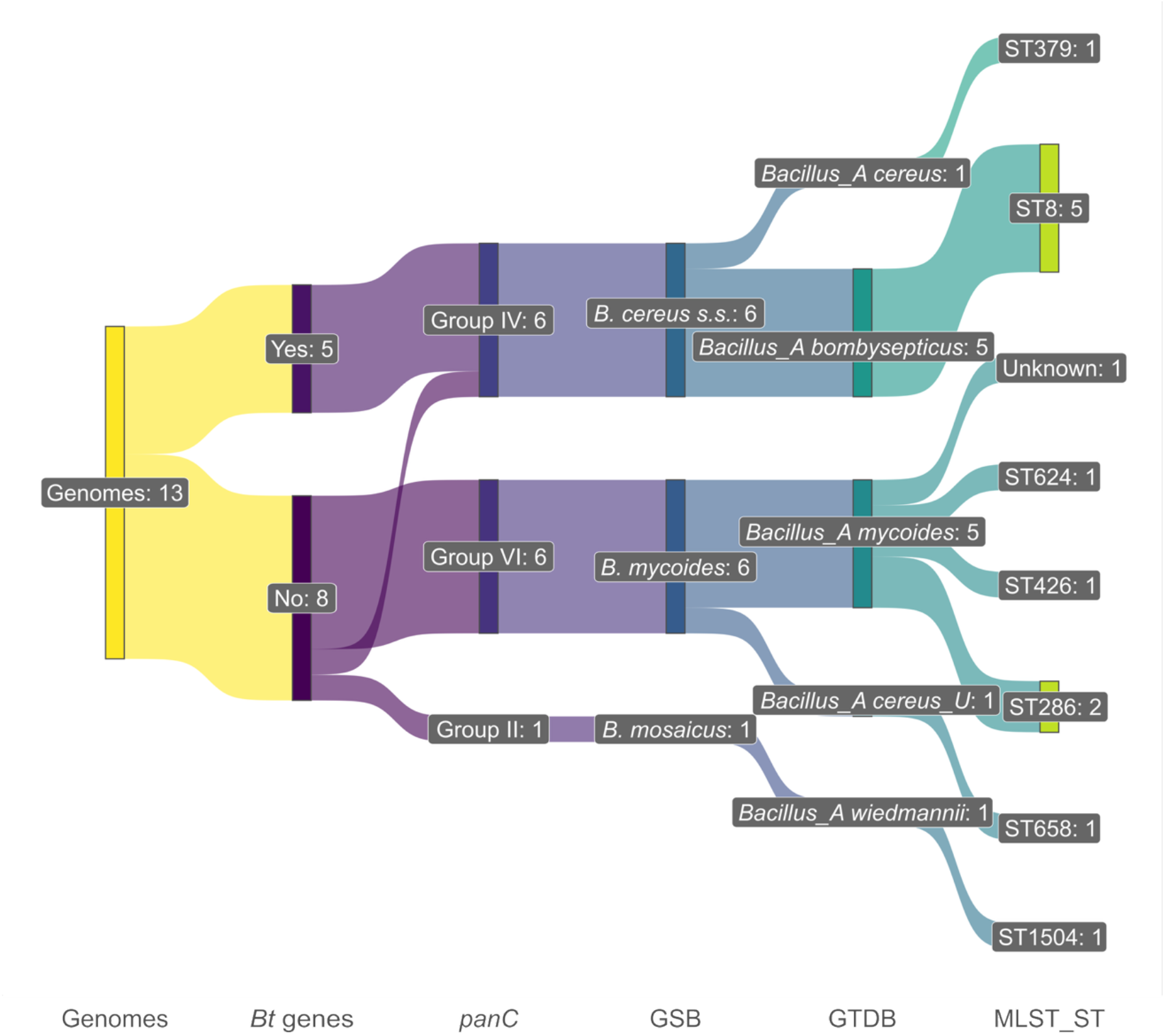
Sankey plot summarising taxonomic assignments of the novel B. cereus group strains sequenced in this study. From left to right: “Bt genes” = presence/absence of Bt-associated toxin genes, as per BTyper3; “panC” = panC Group assigned via BTyper3; “GSB” = species assigned using BTyper3’s 2020 Genomospecies-Subspecies-Biovar (GSB) taxonomy; “GTDB” = species assigned using the Genome Taxonomy Database (GTDB) Toolkit (GTDB-Tk); “MLST_ST” = sequence type (ST) assigned using PubMLST’s seven-gene multi-locus sequence typing (MLST) scheme for “B. cereus”. Alt. text: Graphical representation of the diversity of novel B. cereus group strains, depending on the taxonomic classification used.

Taken together, regardless of the taxonomic nomenclature used, multiple species/*panC* groups/STs were detected among the 13 strains isolated at the sampling sites (Figure 2; Supplementary Table S4). The *panC* group nomenclature is well-known and accepted, while MLST clades within the *B. cereus* group are thought to reflect habitat- and host-associated patterns (Raymond and Bonsall 2013). Therefore, we will use *panC* group and MLST STs when further discussing the 13 novel strains in comparison to a dataset of publicly available *B. cereus* group genomes (*n* = 2890; Carroll *et al*. 2022).

### The *panC* Group VI and II *B. cereus* group strains exhibited varying degrees of similarity to publicly available genomes

Among the six *panC* Group VI strains, there were five STs represented (Supplementary Figure S3). This included ST286, here composed of two novel strains (243886_shin2 and 243890_shin6) and a publicly available genome from soil in Belgium (strain HuA2-4, NCBI RefSeq Assembly accession GCF_000291175.1; Auwera Van der *et al*. 2013), all within >99.9 ANI of one another (Supplementary Table S10). A third novel strain (243888_shin4) was assigned to ST624, alongside two publicly available genomes (Supplementary Figure S3). Notably, the three genomes did not exhibit high genomic similarity (ANI < 98.5), and instead the closest publicly available genome to 243888_shin4 was a German pasteurized milk isolate (ANI < 99.3; NCBI RefSeq Assembly accession GCF_015845385.1) belonging to ST41 (Supplementary Table S10). The remaining *panC* Group VI novel strains were assigned to ST426 (*n* = 1), ST658 (*n* = 1) and an unknown ST (*n* = 1). There were no representatives of these STs among the publicly available genomes, making them singletons (i.e. only one genome per ST; Supplementary Table S11).

The novel *panC* Group II strain (243887_shin3) was assigned to ST1504 (Supplementary Figure S4). This ST consisted of six genomes: our novel strain and five environmental isolates from the United States (Supplementary Table S11). The novel strain shared comparatively high genomic similarity with the environmental isolates (ANI ≈99.6-99.7), despite originating from a different continent (Supplementary Table S12). Overall, the *panC* Group VI and II strains collected in this study varied in terms of their genomic similarity relative to publicly available genomes.

### The majority of *panC* Group IV *B. cereus* group strains belonged to insecticidal ST8

Of the six novel *panC* Group IV strains, one (243885_shin1) was placed within ST379, of which there were no representatives among the high-quality, publicly available genomes (Figure 3A; Supplementary Table S11). Contrastingly, the majority of the novel *panC* Group IV strains (*n* = 5) were assigned to ST8, which also includes the commercial *Bt* strains (Figure 3B; Supplementary Table S11). All ST8 genomes (*n* = 122, including 117 publicly available, high-quality genomes) were assigned to GTDB species *B. bombysepticus* (Supplementary Tables S4 and S11).

**Figure 3.**
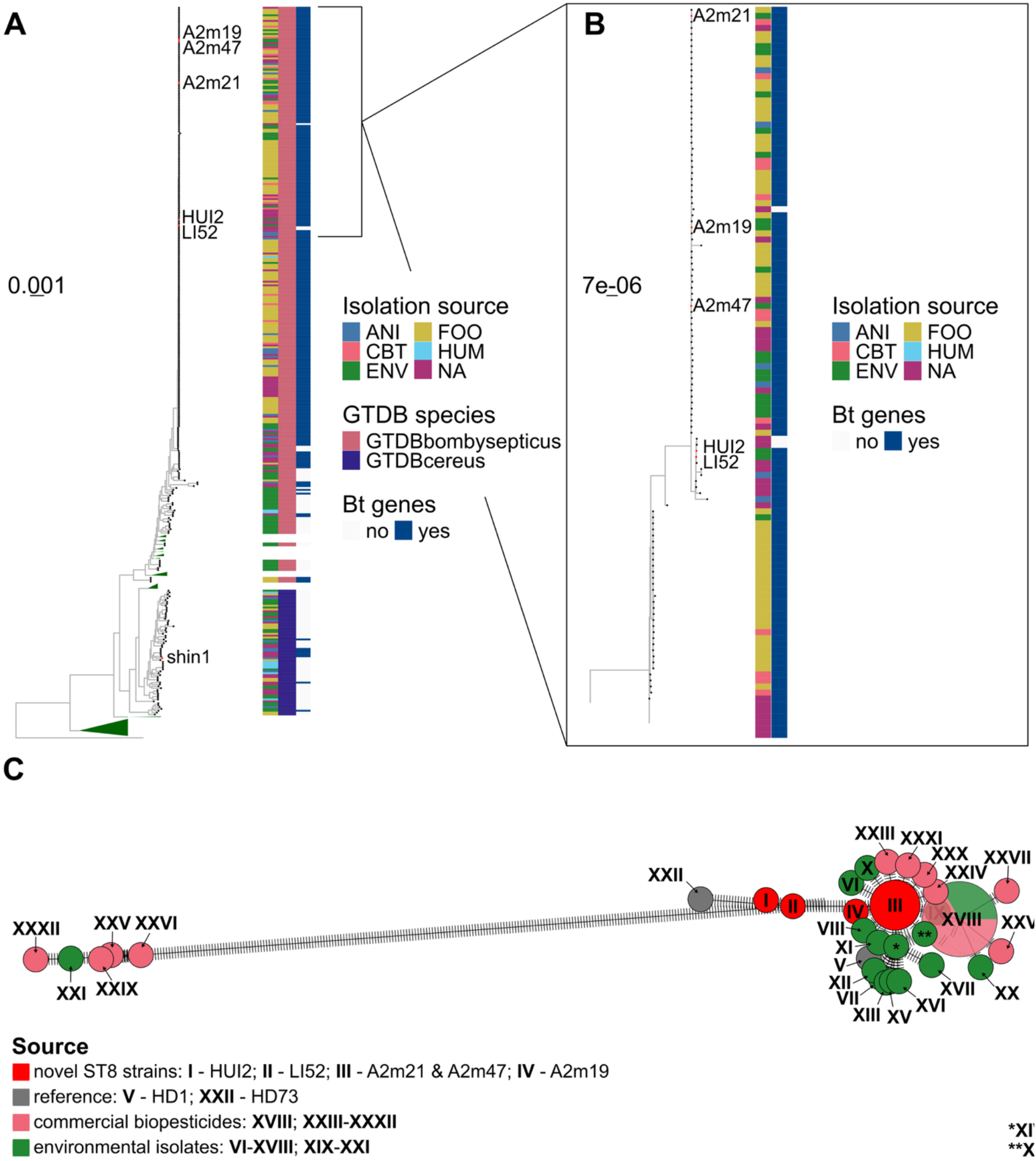
Maximum-likelihood phylogenies of panC Group IV genomes (A) and ST8 genomes (B). Branch lengths represent substitutions per site and novel strains queried in this study are marked with red tips. The numbers in the novel strain names have been omitted for legibility. For (A), clades lacking any of the novel strains have been collapsed (green triangles), and the full tree is available in the supplementary data (Supplementary Figure S6). The phylogeny was rooted using the outgroup (B. anthracis RefSeq Assembly GCF_000008445.1). The ST8 box in panel A includes two genomes not belonging to ST8 (a ST2406 genome and a ST2404 genome), which are not included in panel B. For (B), the phylogeny was rooted using the midpoint. Colour strips to the right of each phylogeny denote (from left to right): (i) isolation source (“Isolation source”), (ii) Genome Taxonomy Database (GTDB) species (“GTDB species”; for Group IV/panel A only), (iii) presence/absence of Bt-associated toxin genes detected by BTyper3 (“Bt genes”). “ANI” = animal-associated isolate; “CBT” = Commercial Bt isolate; “ENV” = environmental isolate; “FOO” = food isolate; “HUM” = human-associated isolate; “NA” = not available. (C) Core SNP haplotype network of environmental, commercial biopesticidal, novel and reference ST8 genomes (Supplementary Table S14). The circles are coloured based on the source, and the size of the circle denotes the number of genomes belonging to a certain haplotype. Alt. text: Graphs illustrating relatedness within the ST8 lineage, in particular how the five novel ST8 strains relate to publicly available ST8 genomes, with subfigures labelled A to C.

We further relied on ANI- and core SNP-based comparisons to examine how closely related the novel strains were to public ST8 biopesticide strains (Supplementary Tables S9 and S13). Three of the novel strains (243882_A2m21, 243884_A2m47 and 243883_A2m19) were within ≥99.9 ANI of public biopesticidal genomes, with core SNP distances ranging between 3-407 SNPs (Supplementary Figure S5). Notably, 243882_A2m21 and 243884_A2m47 were both within ≤5 core SNPs of publicly available biopesticidal strains, and all of the three genomes were within ≤3 core SNPs of NCBI RefSeq Assembly GCF_000835235.1 (an HD-1 *kurstaki* strain isolated from insect larvae) and a *Bt* food isolate from France. 243877_HUI2 shared ≥99.9 ANI with public biopesticidal strains; however, the only ST8 genome within ≤5 core SNPs was 243878_LI52. 243878_LI52 itself was not within 99.9 ANI of any publicly available biopesticidal strains (core SNPs: 334-1813; Supplementary Figure S5). Overall, the novel strains differed by 8-1811 core SNPs from the publicly available environmental ST8 strains, while the equivalent for the public biopesticidal strains was 3-1813 core SNPs (Supplementary Table S9). Thus, the ST8 lineage exhibited a relatively high degree of genomic similarity.

### A higher quantity of *Bt*-associated toxin genes was detected at Bracknell Forest compared to the Orkney Islands

As five out of six novel ST8 strains were positive for *Bt*-associated toxin genes, we conducted further analysis of these insecticidal virulence factors. The five positive genomes split into two groups: a group of two genomes with eight BTyper3 hits for *Bt*-associated toxin genes each (243877_HUI2 and 243878_LI52), and a group of three genomes with 12 BTyper3 hits for *Bt*-associated toxin genes each (243882_A2m21, 243883_A2m19 and 243884_A2m47; Supplementary Table S4; Figure 4). Cumulatively, the number of *Bt*-associated toxin gene hits across all five genomes was 52, with the most prevalent protein families (henceforth referred to as “Cry-families”) being Cry1 (*n* = 43), followed by Vip3 (*n* = 6) and Cry2 (*n* = 3; Figure 4). While all Cry-families were represented at the Bracknell Forest sampling site, only Cry1 genes were detected among novel strains from the Orkney Islands (Figure 4). To corroborate the *in silico* findings, the isolates from the Bracknell Forest and Orkney Islands sampling sites (not including 243879_SET22) were stained for insecticidal proteins. Of the tested strains, only 243881_A1g9 was negative. Hence, the *in silico* and traditional staining methods were in agreement.

**Figure 4.**
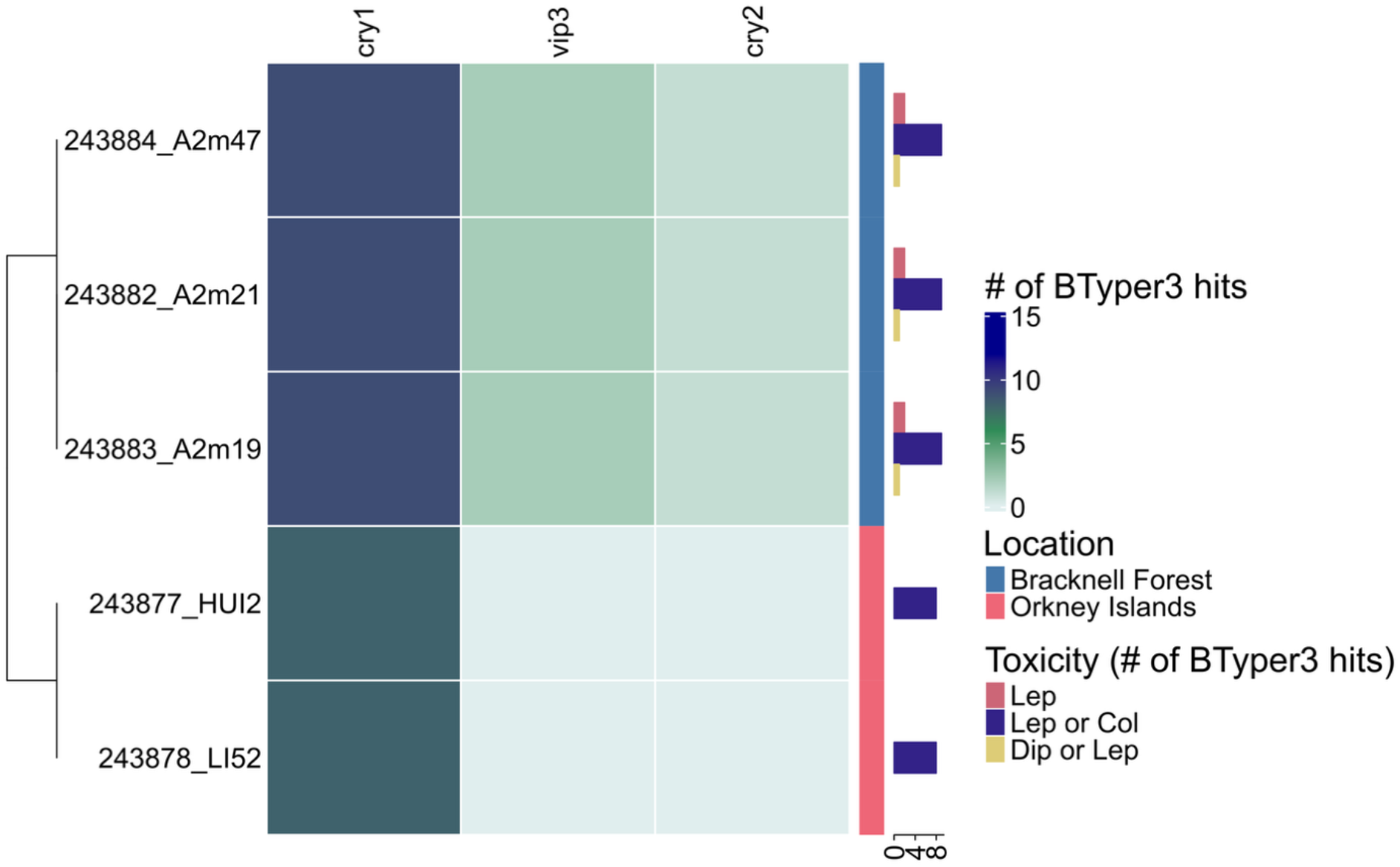
Heatmap of Bt-associated toxin genes detected in novel strains queried in this study (rows), grouped by their respective Cry-families (as per BTyper3 output, columns). Cells are coloured by the number of BTyper3 hits for Bt-associated toxin genes. The colour strip to the right of the heatmap denotes sampling location, while the bar plots show the number of BTyper3 hits per insect target range and genome. The dendrogram represents Euclidean distance based on the Bt-associated toxin matrix. “Lep” = Lepidoptera toxicity; “Lep or Col” = Lepidoptera or Coleoptera toxicity; “Dip or Lep” = Diptera or Lepidoptera toxicity. Alt. text: Graphical representation of the presence/absence of Bt-associated toxin genes among five novel ST8 strains, also illustrating the location of the strains and the toxicity profile of the detected Bt-associated toxin genes.

Cry-families are thought to target different insect orders, and therefore the detected Cry-families were mapped to predicted insect targets using a previously published table (Jouzani, Valijanian, and Sharafi 2017). Briefly, Cry1 proteins are thought to mediate toxicity against Lepidoptera or Coleoptera, Vip3 proteins have been associated with Lepidoptera toxicity and Cry2 proteins have been linked to toxicity against Diptera or Lepidoptera (Jouzani, Valijanian, and Sharafi 2017). Consequently, the most prevalent insect toxicity profile among the novel strains, based on the BTyper3 hits, was Lepidoptera or Coleoptera (*n* = 43), followed by Lepidoptera (*n* = 6) and Diptera or Lepidoptera (*n* = 3; Figure 4).

Efforts were taken to further narrow down the insect target specificity by searching the BPPRC database for the insect species present at the sampling locations in combination with the observed Cry-toxin families, with no success.

### The majority of BGCs detected among *B. cereus* group strains from insect-dense sampling sites are novel

Beyond the *Bt*-associated toxin genes, a microbe’s arsenal of secondary metabolites constitutes a valuable component in interactions within its ecological niche. To assess the biosynthetic potential of the novel strains investigated here, BGCs were mined with the computational tools antiSMASH (Blin *et al*. 2025) and GECCO (Carroll *et al*. 2021). In total, 274 BGCs (hereafter referred to as “the full BGC dataset”) were detected among the 13 novel strains, with antiSMASH reporting a higher number of BGCs (*n* = 196 BGCs) in comparison to GECCO (*n* = 78 BGCs; Supplementary Table S7).

The clustering of unknown BGCs alongside experimentally validated BGCs has previously been used for functional inference and to assess BGC novelty (Doroghazi *et al*. 2014, Navarro-Muñoz *et al*. 2020). In this study, the full BGC dataset was concatenated with publicly available BGCs from the MIBiG database (v4.0; *n* = 2636 BGCs; Zdouc *et al*. 2024) prior to gene cluster family (GCF) delineation (Figure 5A). There were 1722 GCFs in total, of which 1690 GCFs only contained MIBiG BGCs (*n* = 2628 BGCs), eight GCFs contained a mix of BGCs from MIBiG and novel strains (*n* = 112 BGCs), and 24 GCFs uniquely contained BGCs from the novel strains (*n* = 170 BGCs). The majority of the BGCs from the novel strains therefore did not cluster with known MIBiG BGCs (62.04%, rounded to three significant figures), suggesting novelty.

**Figure 5.**
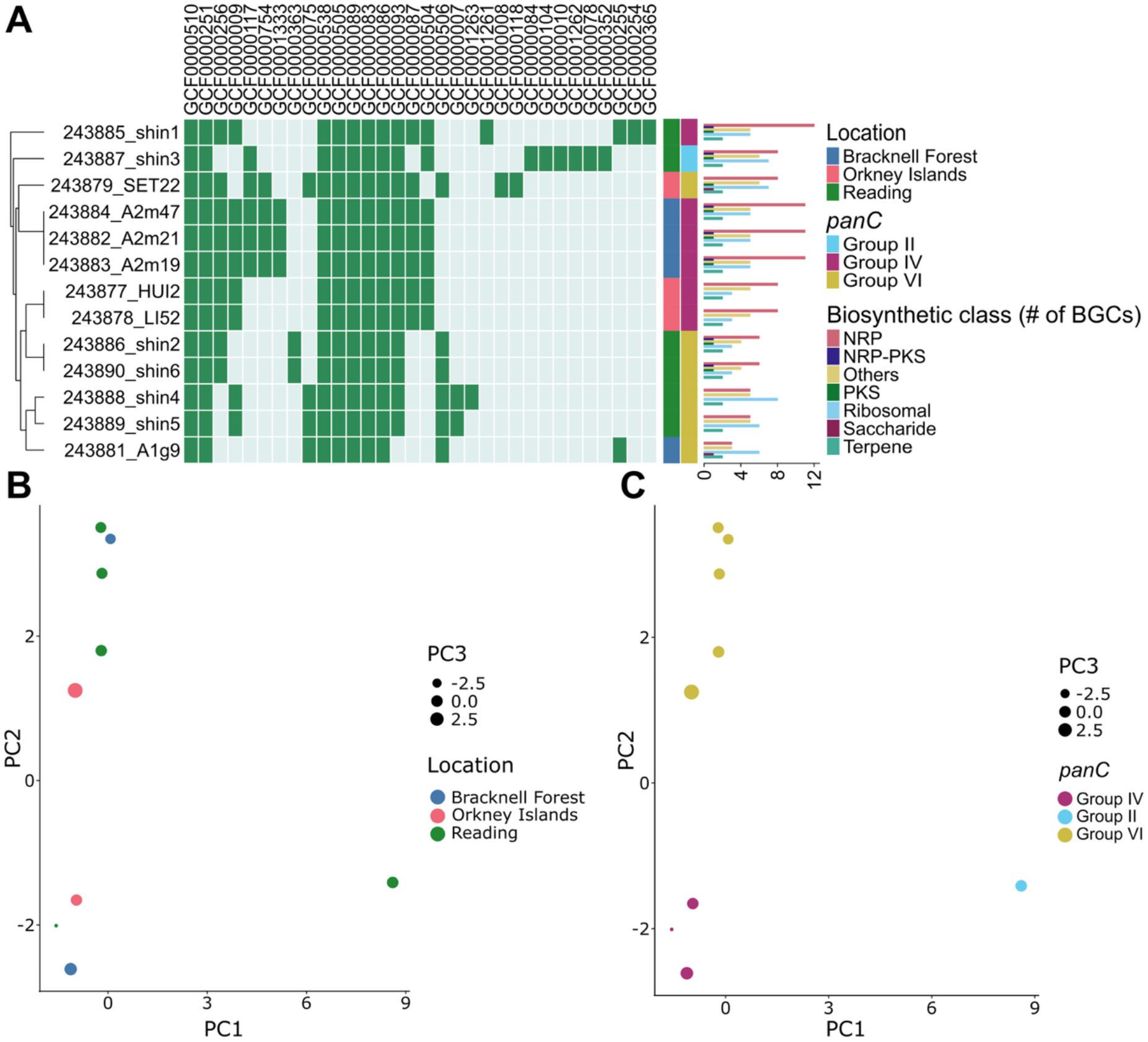
(A) The presence/absence of biosynthetic gene clusters (BGCs) detected among novel strains queried in this study (rows), grouped by gene cluster family (GCFs; columns). Colour strips to the right of the heatmap denote (from left to right): (i) sampling location (“Location”), (ii) panC Group according to BTyper3 (“panC”). Biosynthetic class distributions of the BGCs are shown per genome as bar plots. The dendrogram to the left shows Euclidean distance based on the BGC presence/absence matrix (i.e. the heatmap). “NRP” = non-ribosomally synthesized peptide; “NRP-PKS” = non-ribosomally synthesized peptide-polyketide; “PKS” = polyketide. Principal component analysis (PCA) plots of BGC presence/absence in GCFs for novel strains investigated in this study, annotated based on sampling location (B) or panC Group (according to BTyper3; C). Alt. text: Graphs demonstrating the biosynthetic gene cluster profiles of the 13 novel B. cereus group strains, with subfigures labelled from A to C.

To examine whether the BGC repertoire was linked to environment (i.e. ecological niches) or strain-specific (here, *panC* group assignment), we performed principal component analysis and Euclidean distance-based clustering on the GCF presence/absence matrix for the 13 novel strains (Figure 5). The groupings of the novel strains observed for both the Euclidean distance clustering and the principal component analysis reflected *panC* group assignments, rather than geographical variation dependent on sampling site (Figure 5).

The novel strains 243882_A2m21, 243884_A2m47 and 243883_A2m19 had identical GCF profiles, reflecting the previously observed high degree of genomic similarity among these three genomes (Figures 3 and 5). One of the GCFs unique to 243882_A2m21, 243884_A2m47 and 243883_A2m19 was GCF0001333 (*n* = 13 BGCs; note that several of the antiSMASH BGCs overlap extensively), which contained BGCs from the three novel strains and the MIBiG BGC for the antimicrobial compound thuricin (BGC0000626; Figure 6; Supplementary Table S7; Rea *et al*. 2010). The BGCs from the novel strains showed a marked resemblance to both thuricin and the insecticidal zwittermicin A (BGC0001059; not present in GCF0001333), with the majority of genes having a >90% amino acid identity to the equivalent genes in the MIBiG BGCs. Intriguingly, while the thuricin gene cluster appeared complete, the BGCs from the novel strains lacked the kanosamine gene cluster typically contained within the zwittermicin A BGC (Figure 6), seemingly replaced with a hypothetical protein (based on BLASTp results). Based on genome mining using GECCO and antiSMASH, zwittermicin A was present in several of the publicly available biopesticidal strains, and the kanosamine gene cluster was absent in four of the strains (SRR9600188, SRR9600290, SRR9600276 and SRR9600274; Supplementary Figure S7). The remaining zwittermicin A BGCs were placed on a contig edge, preventing us from confidently inferring if the same genomic structural alteration was present. None of the other novel strains contained any detected BGCs with a high similarity to zwittermicin A (Supplementary Table S7).

**Figure 6.**
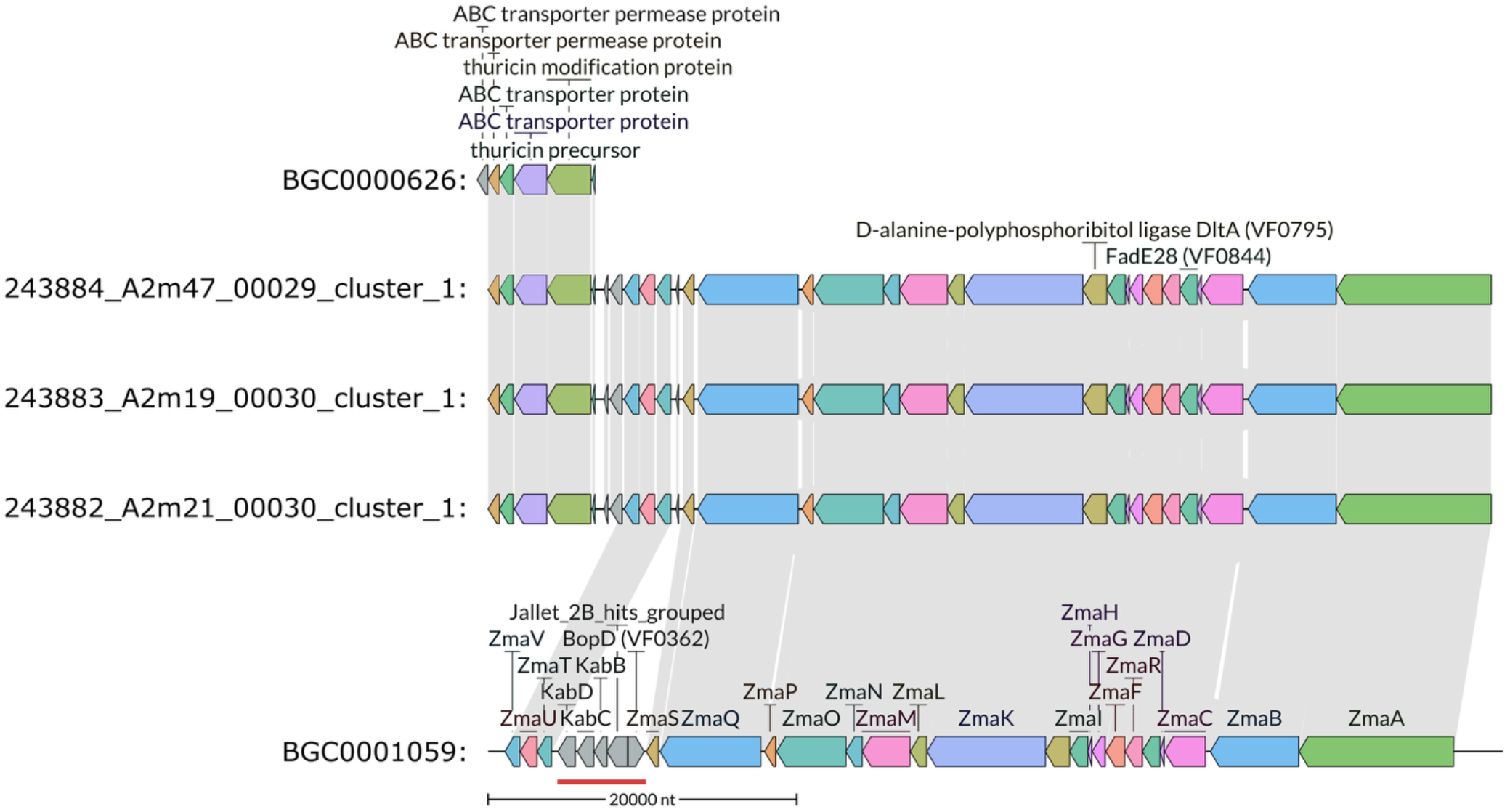
Comparison of GECCO-derived biosynthetic gene clusters (BGCs) present in gene cluster family (GCF) GCF0001333. Shaded links indicate amino acid similarity of 25% with 80% coverage of the shorter sequence within the alignment, while the genes are coloured based on conservation across BGCs. Also included are the MIBiG representatives for thuricin (BGC0000626) and zwittermicin A (BGC0001059). The red bar highlights the kanosamine cluster in the zwittermicin A representative. Alt. text: Graphical representation of biosynthetic gene clusters and their similarity.

## DISCUSSION

### A diverse range of *B. cereus* group strains can be detected at sites with high-density insect populations

The *B. cereus* group species complex encompasses an array of related bacteria, known to inhabit diverse environments and hosts. In this study, we focused on interactions between *B. cereus* group members and insects, aiming to discover putative novel insecticidal compounds with the potential to be used as safe biopesticides. Characterization of isolates from three UK sites with high-density insect populations resulted in a diverse range of *B. cereus* group members, spanning multiple species and eight STs. Other UK-based field trials have previously noted ST8 to be among the most common, if not the dominant, ST in areas with no prior *Bt* application (Bizzarri, Prabhakar, and Bishop 2008, Raymond *et al*. 2010). Similarly, ST8 was the most prevalent ST among our novel strains, supporting the natural occurrence of this ST in UK environments. Intriguingly, one novel strain could not be assigned to an ST. The MLST results regarding this novel strain were inconclusive, and thus we could not determine whether the lack of assignment was due to incomplete or truncated MLST loci not facilitating comprehensive sequence typing, or because the novel strain belonged to a previously undetected ST.

### *Bt*-associated toxin-carrying strains from areas with high-density insect populations were closely related to biopesticidal strains

ST8 encompasses biopesticidal *Bt* strains used for commercial purposes, including the *B. thuringiensis* var. *kurstaki* HD-1 strain that forms the basis for several *Bt* formulations (Raymond and Federici 2017). Here, dedicated ANI- and core SNP-based analysis was used to compare our novel strains to publicly available ST8 genomes, including biopesticidal and environmental strains. Several novel strains exhibited low core SNP distances relative to publicly available biopesticidal strains (*n* = 3 novel strains differing from publicly available biopesticidal strains by ≤6 core SNPs). While these three novel strains share a high degree of genomic similarity with biopesticidal strains, determining whether they are of biopesticidal origin (e.g. biopesticidal strains applied elsewhere and spread to our sampling locations via air or animal carriers, as noted previously; Eskils and Lövgren 1997, Whaley, Anhold, and Schaalje 1998, Swiecicka 2008) or naturally occurring, environmental strains is challenging based on SNP cutoffs alone. A previous study of the clonal *B. cereus* group ST26 lineage noted that strains linked to an emetic foodborne outbreak in New York State could differ by up to seven SNPs, as determined by Snippy (the same SNP calling method used here; Carroll and Wiedmann 2020). Conversely, two strains of *B. anthracis*, another *B. cereus* group clonal lineage, from a case of reoccurring bovine anthrax in Germany differed by three core SNPs, despite being isolated over a decade apart (Braun *et al*. 2022). Consequently, future studies of ST8 evolution and ecology (e.g. estimation of the ST8 evolutionary rate in the wild, modelling of ST8 spread) are needed to improve ST8 source tracking.

### Patterns of Cry-family toxicity observed among the novel strains reflect insect species observed at the sampling sites

*Bt*-associated toxins can confer insecticidal activity to *B. cereus* group strains, with the target species range dependent on which Cry-family the toxins belong to (Bravo *et al*. 2011, Jouzani, Valijanian, and Sharafi 2017). Cry1 toxins have been linked to Lepidoptera toxicity and are thought to be the most common Cry-family (Jouzani, Valijanian, and Sharafi 2017, Zheng *et al*. 2017). All novel strains with predicted *Bt*-associated toxin genes were positive for Cry1, with the Orkney Islands strains exclusively carrying Cry1 genes. Intriguingly, the Orkney Island strains were isolated from sites with high densities of the Lepidopteran winter moth (*Operophtera brumata* (L.); Graham *et al*. 2004). One of the three genomes from the Orkney Islands, 243879_SET22, was not positive for any *Bt*-associated toxin genes, despite the strain being isolated at the same sampling location.

In comparison to the Orkney Islands isolates, the novel strains from the Bracknell Forest location exhibited a more expansive repertoire of *Bt*-associated toxins, including Cry1, Cry2 and Vip3 proteins. An earlier study described a co-occurrence network of Cry1, Cry2, Cry9 and Vip3 proteins linked to Lepidoptera toxicity (Zheng *et al*. 2017), which bears some similarity to the *Bt*-associated toxins detected among the Bracknell Forest novel strains. However, no Cry9 genes were predicted among the Bracknell Forest genomes. The Bracknell Forest sampling location was characterized by Coleopteran insect *Gastriohysa viridula* (Collier, Elliot, and Ellis 2005), and the only detected Cry-family with a target range including Coleopterans was Cry1.

### *B. cereus* group strains in insect-dense environments are predicted to produce novel, strain-specific BGCs

The novel strains investigated in this study were also mined for BGCs, and a high proportion of the detected BGCs were seemingly novel, suggesting that there could be previously unexplored candidates for biopesticide development. Additionally, three of the novel strains were positive for the insecticidal secondary metabolite zwittermicin A, indicating potential for insecticidal activity. The zwittermicin A BGCs detected among the novel strains exhibited minor differences in comparison to the MIBiG representative BGC for zwittermicin A (originally isolated in “*B. cereus*”; Emmert *et al*. 2004, Kevany, Rasko, and Thomas 2009), as the kanosamine BGC had been lost. This could indicate a streamlining of the genome and adaptation of a defensive repertoire, further supported by the nearby presence of the antimicrobial thuricin BGC. Otherwise, the included genes showed high sequence similarity to the canonical MIBiG BGCs, and therefore it is likely that the produced zwittermicin A compound will exhibit little variation to previously isolated examples. Similar genomic streamlining was observed for four of the public biopesticidal strains. Contrastingly, the remaining two novel ST8 strains were negative for zwittermicin A, which was expected given the close genomic similarity to the zwittermicin A-negative HD-73 strain (Zhu *et al*. 2015).

The clustering of the novel strains based on GCF presence/absence reflected species (here, *panC* groups) rather than location-based patterns. Hence, it appears that, despite *B. cereus* group members being highly genetically related, species boundaries were sufficient to prevent BGC transfer among our novel strains. However, this study is limited in terms of the number of isolates, sampling locations and represented *panC* groups, and future research with larger samples could therefore result in different findings.

### *B. cereus* group strains from the insect cadaver: toxin cheats or insecticidal secondary metabolite producers?

The classical *Bt* strains are known to be obligate killers, where inducing mortality in the insect host is required for reproduction (Raymond and Erdos 2022). The isolation of *B. cereus* group strains from insect cadavers has previously sourced potent, biopesticidal strains (Dulmage 1970, Barjac de 1978), yet epizootics and even putative *Bt-*killed cadavers are rare in the field (Porcar and Caballero 2000, Konecka *et al*. 2007, Alejandro López-Pazos *et al*. 2009, Khojand *et al*. 2013). Therefore, the novel strains from the insect cadaver in Reading are of particular interest; however, none of the novel strains from the dead insect were positive for *Bt*-associated toxin genes, for which we here consider two potential explanations. Firstly, it has been noted that mixed *B. cereus* group infections can occur in insects, where a proportion of the infecting bacteria lack *Bt*-associated toxins (Raymond and Bonsall 2013, Raymond and Erdos 2022). The toxin-lacking strains can reproduce in the insect host by benefiting from public goods, i.e. the toxin secreted by other bacteria. As the production of *Bt*-associated toxins is metabolically expensive, the toxin-lacking strains may outcompete the toxin-carrying strains in mixed infections (Raymond and Bonsall 2013, Raymond and Erdos 2022). It is therefore plausible that the strains isolated from the insect cadaver were “toxin cheats”, and thus not the causative agents of the insect’s demise. Competition experiments conducted at temperatures that are field-realistic for the UK (15 °C) show that *B. mycoides* isolates can readily outcompete a *Bt* Cry toxin producer in insect cadavers (Manktelow *et al*. 2021). However, intense sampling of insect cadavers followed by WGS, and hence genomic evidence of mixed infections, is very rare.

A second explanation is that the insect was killed by entomopathogenic secondary metabolites produced by one or several of the isolated strains. Predicted BGCs detected among the novel strains queried here showed an overall high degree of novelty, and one of the strains from the dead insect (243885_shin1) carried the highest number of BGCs present in novel GCFs (i.e. GCFs lacking MIBiG BGCs). Further investigation of the novel secondary metabolites produced by 243885_shin1 may generate prospective insecticidal compound leads, which could be developed for industrial applications.

## CONCLUSION

In this investigation, we aimed to provide an overview of *B. cereus* group-insect interactions by examining the epidemiology, *Bt*-associated toxins and BGCs of 13 novel *B. cereus* group strains isolated from regions of the UK with high-density insect populations, but no prior history of commercial *Bt* application. The results demonstrated a high diversity of *B. cereus* group members, as well as confirmed the presence of *Bt*-associated toxin genes relevant to insect targets in close proximity. We also identified novel BGCs with unknown bioactivities, in particular among *B. cereus* group strains isolated from an insect cadaver. The impact of this study is limited by the comparatively small sample size, and future studies with expanded sample sizes are needed. However, the findings represent a first proof-of-concept of genomic mining of *B. cereus* group strains from sites with high-density insect populations with the aim of generating novel compound leads for biopesticide development.

## AUTHOR CONTRIBUTIONS

JB - Formal analysis, Investigation, Visualisation, Writing - original draft, Writing-review and editing; VR – Formal analysis, Writing-review and editing; BR - Conceptualization, Funding acquisition, Project administration, Resources, Visualisation, Writing-review and editing; LMC – Conceptualization, Funding acquisition, Project administration, Supervision, Resources, Writing-review and editing.

## REFERENCES

Alejandro López-Pazos S, Wilson Martínez J, Castillo AX et al. Presence and significance of *Bacillus thuringiensis* Cry proteins associated with the Andean weevil *Premnotrypes vorax* (Coleoptera: Curculionidae). Rev Biol Trop 2009;57:1235–43. DOI: 10.15517/rbt.v57i4.5460.

Auwera GA Van der, Feldgarden M, Kolter R et al. Whole-genome sequences of 94 environmental isolates of *Bacillus cereus sensu lato*. Genome Announc 2013;1: e00380–13. DOI: 10.1128/genomea.00380-13.

Barjac H de. Une nouvelle variété de *Bacillus thuringiensis* très toxique pour les moustiques: *B. thuringiensis* var. *israelensis* sérotype 14 [A new variety of *Bacillus thuringinesis* very toxic to mosquitoes: *B. thuringiensis* var. *israelensis* serotype 14]. C R Acad Hebd Seances Acad Sci D 1978;286:797–800.

Bizzarri MF, Prabhakar A, Bishop AH. Multiple-locus sequence typing analysis of *Bacillus thuringiensis* recovered from the phylloplane of clover (*Trifolium hybridum*) in vegetative form. Microb Ecol 2008;55:619–25. DOI: 10.1007/s00248-007-9305-3.

Blin K, Shaw S, Steinke K et al. AntiSMASH 5.0: updates to the secondary metabolite genome mining pipeline. Nucleic Acids Res 2019;47:W81–7. DOI: 10.1093/nar/gkz310.

Blin K, Shaw S, Vader L et al. AntiSMASH 8.0: extended gene cluster detection capabilities and analyses of chemistry, enzymology, and regulation. Nucleic Acids Res 2025;53:W32–8. DOI: 10.1093/nar/gkaf334.

Blom J, Wambui J, Gourlé H et al. Large-scale insights into the biosynthetic potential of the *Bacillus cereus* group. bioRxiv 2025. DOI: 10.1101/2025.10.16.682773.

Bode HB. Entomopathogenic bacteria as a source of secondary metabolites. Curr Opin Chem Biol 2009;13:224–30. DOI: 10.1016/j.cbpa.2009.02.037.

Braun P, Beyer W, Hanczaruk M et al. Reoccurring bovine anthrax in Germany on the same pasture after 12 years. J Clin Microbiol 2022;60:e02291–21. DOI: 10.1128/jcm.02291-21.

Bravo A, Gómez I, Porta H et al. Evolution of *Bacillus thuringiensis* Cry toxins insecticidal activity. Microb Biotechnol 2013;6:17–26. DOI: 10.1111/j.1751-7915.2012.00342.x.

Bravo A, Likitvivatanavong S, Gill SS et al. *Bacillus thuringiensis*: a story of a successful bioinsecticide. Insect Biochem Mol Biol 2011;41:423–31. DOI: 10.1016/j.ibmb.2011.02.006.

Carroll LM, Cheng RA, Kovac J. No assembly required: using BTyper3 to assess the congruency of a proposed taxonomic framework for the *Bacillus cereus* group with historical typing methods. Front Microbiol 2020;11:580691. DOI: 10.3389/fmicb.2020.580691.

Carroll LM, Huisman JS, Wiedmann M. Twentieth-century emergence of antimicrobial resistant human- and bovine-associated *Salmonella enterica* serotype Typhimurium lineages in New York State. Sci Rep 2020;10:14428. DOI: 10.1038/s41598-020-71344-9.

Carroll LM, Larralde M, Fleck JS et al. Accurate *de novo* identification of biosynthetic gene clusters with GECCO. bioRxiv 2021. DOI: 10.1101/2021.05.03.442509.

Carroll LM, Marston CK, Kolton CB et al. Strains associated with two 2020 welder anthrax cases in the United States belong to separate lineages within *Bacillus cereus sensu lato*. Pathogens 2022;11:856. DOI: 10.3390/pathogens11080856.

Carroll LM, Wiedmann M. Cereulide synthetase acquisition and loss events within the evolutionary history of group III *Bacillus cereus sensu lato* facilitate the transition between emetic and diarrheal foodborne pathogens. mBio 2020;11: e01263–20. DOI: 10.1128/mBio.01263-20.

Carroll LM, Wiedmann M, Kovac J. Proposal of a taxonomic nomenclature for the *Bacillus cereus* group which reconciles genomic definitions of bacterial species with clinical and industrial phenotypes. mBio 2020;11:e00034–20. DOI: 10.1128/mBio.

Chaumeil PA, Mussig AJ, Hugenholtz P et al. GTDB-Tk: a toolkit to classify genomes with the Genome Taxonomy Database. Bioinformatics 2020;36:1925–7. DOI: 10.1093/bioinformatics/btz848.

Chen S, Zhou Y, Chen Y et al. Fastp: An ultra-fast all-in-one FASTQ preprocessor. Bioinformatics 2018;34:i884–90. DOI: 10.1093/bioinformatics/bty560.

Collier FA, Elliot SL, Ellis RJ. Spatial variation in *Bacillus thuringiensis/cereus* populations within the phyllosphere of broad-leaved dock (*Rumex obtusifolius*) and surrounding habitats. FEMS Microbiol Ecol 2005;54:417–25. DOI: 10.1016/j.femsec.2005.05.005.

Crickmore N, Berry C, Panneerselvam S et al. A structure-based nomenclature for *Bacillus thuringiensis* and other bacteria-derived pesticidal proteins. J Invertebr Pathol 2021;186:107438. DOI: 10.1016/j.jip.2020.107438.

Croucher NJ, Page AJ, Connor TR et al. Rapid phylogenetic analysis of large samples of recombinant bacterial whole genome sequences using Gubbins. Nucleic Acids Res 2015;43:e15. DOI: 10.1093/nar/gku1196.

Day M, Ibrahim M, Dyer D et al. Genome sequence of *Bacillus thuringiensis* subsp. *Kurstaki* strain HD-1. Genome Announc 2014;2:e00613-14. DOI: 10.1128/genomeA.00613-14.

Doroghazi JR, Albright JC, Goering AW et al. A roadmap for natural product discovery based on large-scale genomics and metabolomics. Nat Chem Biol 2014;10:963–8. DOI: 10.1038/nchembio.1659.

Dulmage HT. Insecticidal activity of HD-1, a new isolate *Bacillus thuringiensis* var. *alesti*. J Invertebr Pathol 1970;15:232–9. DOI: 10.1016/0022-2011(70)90240-5.

Egorov AA, Atkinson GC. LoVis4u: a locus visualization tool for comparative genomics and coverage profiles. NAR Genom Bioinform 2025;7:lqaf009. DOI: 10.1093/nargab/lqaf009.

Ehling-Schulz M, Frenzel E, Gohar M. Food-bacteria interplay: pathometabolism of emetic *Bacillus cereus*. Front Microbiol 2015;6:704. DOI: 10.3389/fmicb.2015.00704.

Emmert EAB, Klimowicz AK, Thomas MG et al. Genetics of zwittermicin A production by *Bacillus cereus*. Appl Environ Microbiol 2004;70:104–13. DOI: 10.1128/AEM.70.1.104-113.2004.

Eskils K, Lövgren A. Release of *Bacillus thuringiensis* subsp. *israelensis* in Swedish soil. FEMS Microbiol Ecol 1997;23:229–37. DOI: 10.1111/j.1574-6941.1997.tb00405.x.

Ewels P, Magnusson M, Lundin S et al. MultiQC: summarize analysis results for multiple tools and samples in a single report. Bioinformatics 2016;32:3047–8. DOI: 10.1093/bioinformatics/btw354.

Feldgarden M, Brover V, Haft DH et al. Validating the AMRFinder tool and resistance gene database by using antimicrobial resistance genotype-phenotype correlations in a collection of isolates. Antimicrob Agents Chemother 2019;63:e00483–19. DOI: 10.1128/AAC.00483-19.

Galili T. dendextend: an R package for visualizing, adjusting and comparing trees of hierarchical clustering. Bioinformatics 2015;31:3718–20. DOI: 10.1093/bioinformatics/btv428.

Gilchrist CLM, Chooi YH. clinker & clustermap.js: automatic generation of gene cluster comparison figures. Bioinformatics 2021;37:2473–5. DOI: 10.1093/bioinformatics/btab007.

Graham RI, Tyne WI, Possee RD et al. Genetically variable nucleopolyhedroviruses isolated from spatially separate populations of the winter moth *Operophtera brumata* (Lepidoptera: Geometridae) in Orkney. J Invertebr Pathol 2004;87:29–38. DOI: 10.1016/j.jip.2004.06.002.

Grubbs KJ, Bleich RM, Santa Maria KC et al. Large-scale bioinformatics analysis of *Bacillus* genomes uncovers conserved roles of natural products in bacterial physiology. mSystems 2017;2:e00040–17. DOI: 10.1128/mSystems.00040-17.

Gu Z, Eils R, Schlesner M. Complex heatmaps reveal patterns and correlations in multidimensional genomic data. Bioinformatics 2016;32:2847–9. DOI: 10.1093/bioinformatics/btw313.

Gu Z, Gu L, Eils R et al. circlize implements and enhances circular visualization in R. Bioinformatics 2014;30:2811–2. DOI: 10.1093/bioinformatics/btu393.

Gurevich A, Saveliev V, Vyahhi N et al. QUAST: quality assessment tool for genome assemblies. Bioinformatics 2013;29:1072–5. DOI: 10.1093/bioinformatics/btt086.

Jain C, Rodriguez-R LM, Phillippy AM et al. High throughput ANI analysis of 90K prokaryotic genomes reveals clear species boundaries. Nat Commun 2018;9:5114. DOI: 10.1038/s41467-018-07641-9.

Jolley KA, Maiden MCJ. BIGSdb: Scalable analysis of bacterial genome variation at the population level. BMC Bioinformatics 2010;11:595. DOI: 10.1186/1471-2105-11-595.

Jouzani GS, Valijanian E, Sharafi R. *Bacillus thuringiensis*: a successful insecticide with new environmental features and tidings. Appl Microbiol Biotechnol 2017;101:2691– 711. DOI: 10.1007/s00253-017-8175-y.

Kalyaanamoorthy S, Minh BQ, Wong TKF et al. ModelFinder: fast model selection for accurate phylogenetic estimates. Nat Methods 2017;14:587–9. DOI: 10.1038/nmeth.4285.

Kevany BM, Rasko DA, Thomas MG. Characterization of the complete zwittermicin A biosynthesis gene cluster from *Bacillus cereus*. Appl Environ Microbiol 2009;75:1144–55. DOI: 10.1128/AEM.02518-08.

Khojand S, Keshavarzi M, Zargari K et al. Presence of multiple cry genes in *Bacillus thuringiensis* isolated from dead cotton bollworm *Heliothis armigera*. J Agr Sci Tech 2013;15:1285–92.

Konecka E, Kaznowski A, Ziemnicka J et al. Molecular and phenotypic characterisation of *Bacillus thuringiensis* isolated during epizootics in *Cydia pomonella* L. J Invertebr Pathol 2007;94:56–63. DOI: 10.1016/j.jip.2006.08.008.

la Fuente-Salcido NM de, Casados-Vázquez LE, Barboza-Corona JE. Bacteriocins of *Bacillus thuringiensis* can expand the potential of this bacterium to other areas rather than limit its use only as microbial insecticide. Can J Microbiol 2013;59:515–22. DOI: 10.1139/cjm-2013-0284.

Larralde M, Blom J, Gourlé H et al. Fast, flexible gene cluster family delineation with IGUA. bioRxiv 2025. DOI: 10.1101/2025.05.15.654203.

Levinson BL, Kasyan KJ, Chiu SS et al. Identification of beta-exotoxin production, plasmids encoding beta-exotoxin, and a new exotoxin in *Bacillus thuringiensis* by using high-performance liquid chromatography. J Bacteriol 1990;172:3172–9. DOI: 10.1128/jb.172.6.3172-3179.1990.

Liu G, Song L, Shu C et al. Complete genome sequence of *Bacillus thuringiensis* subsp. *kurstaki* strain HD73. Genome Announc 2013;1:e00080–13. DOI: 10.1128/genomeA.00080-13.

Liu XY, Ruan LF, Hu ZF et al. Genome-wide screening reveals the genetic determinants of an antibiotic insecticide in *Bacillus thuringiensis*. J Biol Chem 2010;285:39191–200. DOI: 10.1074/jbc.M110.148387.

Maagd RA de, Bravo A, Crickmore N. How *Bacillus thuringiensis* has evolved specific toxins to colonize the insect world. Trends Genet 2001;17:193–9. DOI: 10.1016/s0168-9525(01)02237-5.

Manktelow CJ, White H, Crickmore N et al. Divergence in environmental adaptation between terrestrial clades of the *Bacillus cereus* group. FEMS Microbiol Ecol 2021;97:fiaa228. DOI: 10.1093/femsec/fiaa228.

Martínez C, Porcar M, López A et al. Characterization of a *Bacillus thuringiensis* strain with a broad spectrum of activity against lepidopteran insects. Entomol Exp Appl 2004;111:71–7. DOI: 10.1111/j.0013-8703.2004.00156.x.

Minh BQ, Schmidt HA, Chernomor O et al. IQ-TREE 2: new models and efficient methods for phylogenetic inference in the genomic era. Mol Biol Evol 2020;37:1530–4. DOI: 10.1093/molbev/msaa015.

Mondol MAM, Shin HJ, Islam MT. Diversity of secondary metabolites from marine *Bacillus* species: chemistry and biological activity. Mar Drugs 2013;11:2846–72. DOI: 10.3390/md11082846.

Mullowney MW, Duncan KR, Elsayed SS et al. Artificial intelligence for natural product drug discovery. Nat Rev Drug Discov 2023;22:895–916. DOI: 10.1038/s41573-023-00774-7.

Navarro-Muñoz JC, Selem-Mojica N, Mullowney MW et al. A computational framework to explore large-scale biosynthetic diversity. Nat Chem Biol 2020;16:60–8. DOI: 10.1038/s41589-019-0400-9.

Page AJ, Taylor B, Delaney AJ et al. SNP-sites: rapid efficient extraction of SNPs from multi-FASTA alignments. Microb Genom 2016;2:e000056. DOI: 10.1099/mgen.0.000056.

Paradis E. pegas: an R package for population genetics with an integrated-modular approach. Bioinformatics 2010;26:419–20. DOI: 10.1093/bioinformatics/btp696.

Paradis E, Schliep K. ape 5.0: an environment for modern phylogenetics and evolutionary analyses in R. Bioinformatics 2019;35:526–8. DOI: 10.1093/bioinformatics/bty633.

Parks DH, Imelfort M, Skennerton CT et al. CheckM: assessing the quality of microbial genomes recovered from isolates, single cells, and metagenomes. Genome Res 2015;25:1043–55. DOI: 10.1101/gr.186072.114.

Petit III RA, Read TD. Bactopia: a flexible pipeline for complete analysis of bacterial genomes. mSystems 2020;5:e00190–20. DOI: 10.1128/msystems.00190-20.

Porcar M, Caballero P. Molecular and insecticidal characterization of a *Bacillus thuringiensis* strain isolated during a natural epizootic. J Appl Microbiol 2000;89:309–16. DOI: 10.1046/j.1365-2672.2000.01115.x.

Ragasruthi M, Balakrishnan N, Murugan M et al. *Bacillus thuringiensis* (Bt)-based biopesticide: navigating success, challenges, and future horizons in sustainable pest control. Sci Total Environ 2024;954:176594. DOI: 10.1016/j.scitotenv.2024.176594.

Raymond B, Bonsall MB. Cooperation and the evolutionary ecology of bacterial virulence: the *Bacillus cereus* group as a novel study system. BioEssays 2013;35:706–16. DOI: 10.1002/bies.201300028.

Raymond B, Erdos Z. Passage and the evolution of virulence in invertebrate pathogens: fundamental and applied perspectives. J Invertebr Pathol 2022;187:107692. DOI: 10.1016/j.jip.2021.107692.

Raymond B, Federici BA. In defense of *Bacillus thuringiensis*, the safest and most successful microbial insecticide available to humanity - a response to EFSA. FEMS Microbiol Ecol 2017;93:fix084. DOI: 10.1093/femsec/fix084.

Raymond B, Wyres KL, Sheppard SK et al. Environmental factors determining the epidemiology and population genetic structure of the *Bacillus cereus* group in the field. PLoS Pathog 2010;6:e1000905. DOI: 10.1371/journal.ppat.1000905.

Rea MC, Sit CS, Clayton E et al. Thuricin CD, a posttranslationally modified bacteriocin with a narrow spectrum of activity against *Clostridium difficile*. Proc Natl Acad Sci U S A 2010;107:9352–7. DOI: 10.1073/pnas.0913554107.

Revell LJ. phytools 2.0: an updated R ecosystem for phylogenetic comparative methods (and other things). PeerJ 2024;12:e16505. DOI: 10.7717/peerj.16505.

Schliep KP. phangorn: phylogenetic analysis in R. Bioinformatics 2011;27:592–3. DOI: 10.1093/bioinformatics/btq706.

Seemann T. Prokka: rapid prokaryotic genome annotation. Bioinformatics 2014;30:2068–9. DOI: 10.1093/bioinformatics/btu153.

Song L, Florea L, Langmead B. Lighter: fast and memory-efficient sequencing error correction without counting. Genome Biol 2014;15:509. DOI: 10.1186/s13059-014-0509-9.

Souvorov A, Agarwala R, Lipman DJ. SKESA: strategic k-mer extension for scrupulous assemblies. Genome Biol 2018;19:153. DOI: 10.1186/s13059-018-1540-z.

Staley JT, Stewart-Jones A, Pope TW et al. Varying responses of insect herbivores to altered plant chemistry under organic and conventional treatments. Proc Biol Sci 2010;277:779–86. DOI: 10.1098/rspb.2009.1631.

Stenfors Arnesen LP, Fagerlund A, Granum PE. From soil to gut: *Bacillus cereus* and its food poisoning toxins. FEMS Microbiol Rev 2008;32:579–606. DOI: 10.1111/j.1574-6976.2008.00112.x.

Swiecicka I. Natural occurrence of *Bacillus thuringiensis* and *Bacillus cereus* in eukaryotic organisms: a case for symbiosis. Biocontrol Sci Technol 2008;18:221–39. DOI: 10.1080/09583150801942334.

Tavarc S. Some Probabilistic and Statistical Problems in the Analysis of DNA Sequences. In: Miura, RM (eds.). Some Mathematical Questions in Biology – DNA Sequence Analysis. American Mathematical Society, 1986.

Thi Hoang D, Chernomor O, Haeseler A von et al. UFBoot2: improving the ultrafast bootstrap approximation. Mol Biol Evol 2017;35:518–22. DOI: 10.1093/molbev/msx281.

Tommaso P Di, Chatzou M, Floden EW et al. Nextflow enables reproducible computational workflows. Nat Biotechnol 2017;35:316–9. DOI: 10.1038/nbt.3820.

Tonkin-Hill G, MacAlasdair N, Ruis C et al. Producing polished prokaryotic pangenomes with the Panaroo pipeline. Genome Biol 2020;21:180. DOI: 10.1186/s13059-020-02090-4.

Tyc O, Song C, Dickschat JS et al. The ecological role of volatile and soluble secondary metabolites produced by soil bacteria. Trends Microbiol 2017;25:280–92. DOI: 10.1016/j.tim.2016.12.002.

Vater J, Tam LTT, Jähne J et al. Plant-associated representatives of the *Bacillus cereus* group are a rich source of antimicrobial compounds. Microorganisms 2023;11:2677. DOI: 10.3390/microorganisms11112677.

Whaley WH, Anhold J, Schaalje GB. Canyon drift and dispersion of *Bacillus thuringiensis* and its effects on select nontarget Lepidopterans in Utah. Environ Entomol 1998;27:539–48.

Wickham H, Averick M, Bryan J et al. Welcome to the Tidyverse. J Open Source Softw 2019;4:1686. DOI: 10.21105/joss.01686.

Xia L, Miao Y, Cao A et al. Biosynthetic gene cluster profiling predicts the positive association between antagonism and phylogeny in *Bacillus*. Nat Commun 2022;13:1023. DOI: 10.1038/s41467-022-28668-z.

Yang Z. A space-time process model for the evolution of DNA sequences. Genetics 1995;139:993–1005. DOI: 10.1093/genetics/139.2.993.

Yin QJ, Ying TT, Zhou ZY et al. Species-specificity of the secondary biosynthetic potential in *Bacillus*. Front Microbiol 2023;14:1271418. DOI: 10.3389/fmicb.2023.1271418.

Yu G, Smith DK, Zhu H et al. ggtree: an R package for visualization and annotation of phylogenetic trees with their covariates and other associated data. Methods Ecol Evol 2017;8:28–36. DOI: 10.1111/2041-210X.12628.

Zdouc MM, Blin K, Louwen NLL et al. MIBiG 4.0: advancing biosynthetic gene cluster curation through global collaboration. Nucleic Acids Res 2024;53:D678–90. DOI: 10.1093/nar/gkae1115.

Zhao X, Kuipers OP. Identification and classification of known and putative antimicrobial compounds produced by a wide variety of *Bacillales* species. BMC Genomics 2016;17:882. DOI: 10.1186/s12864-016-3224-y.

Zheng J, Gao Q, Liu L et al. Comparative genomics of *Bacillus thuringiensis* reveals a path to specialized exploitation of multiple invertebrate hosts. mBio 2017;8:e00822–17. DOI: 10.1128/mBio.00822-17.

Zhu L, Peng D, Wang Y et al. Genomic and transcriptomic insights into the efficient entomopathogenicity of *Bacillus thuringiensis*. Sci Rep 2015;5:14129. DOI: 10.1038/srep14129.

