## Supplementary figures and images for "THE INSECTICIDAL POTENTIAL OF *BACILLUS CEREUS* GROUP STRAINS FROM INSECT-DENSE REGIONS OF THE UNITED KINGDOM"

### Supplementary Figure S1

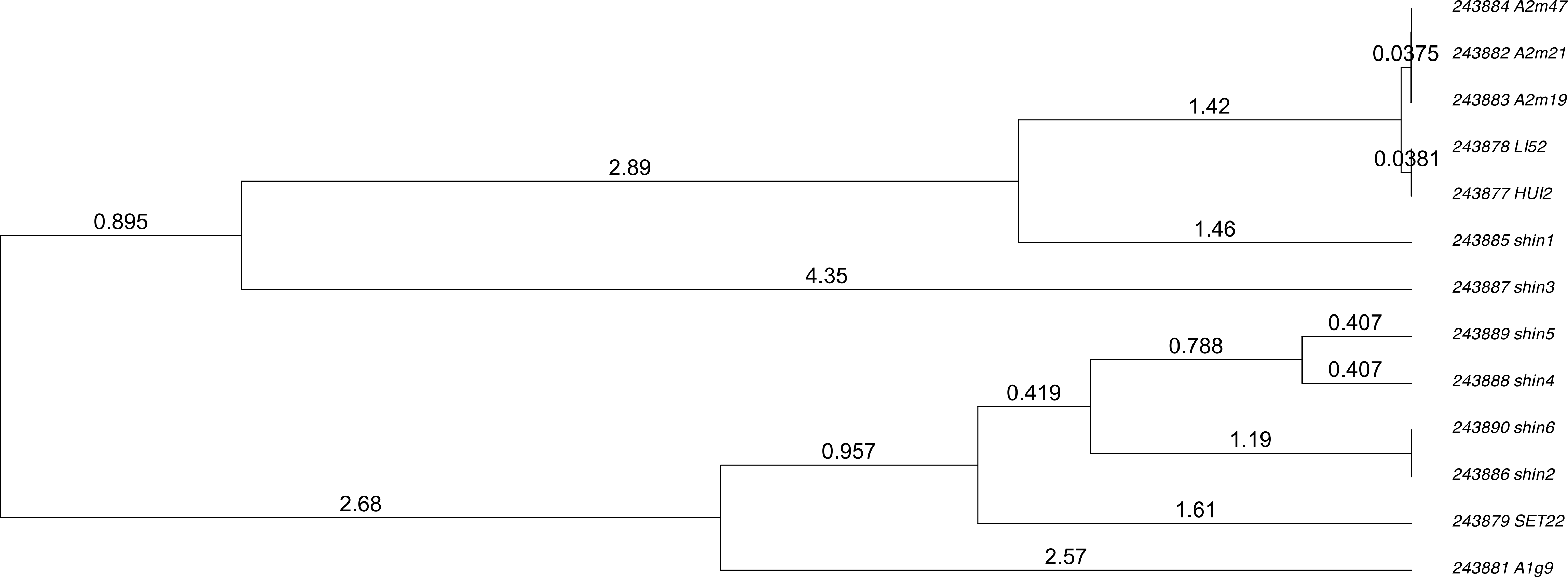

### Supplementary Figure S2

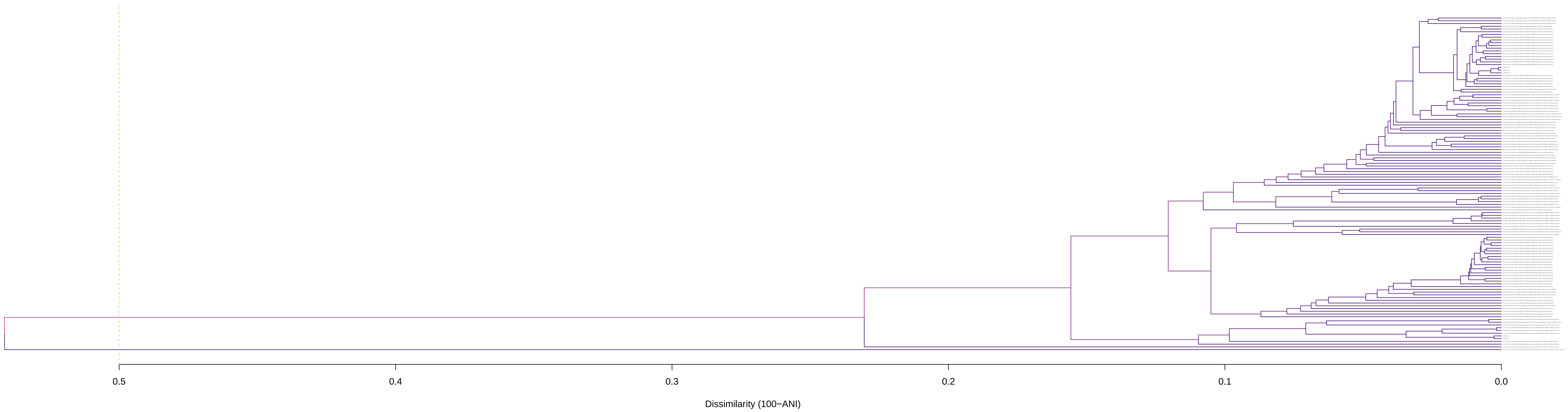

### Supplementary Figure S3

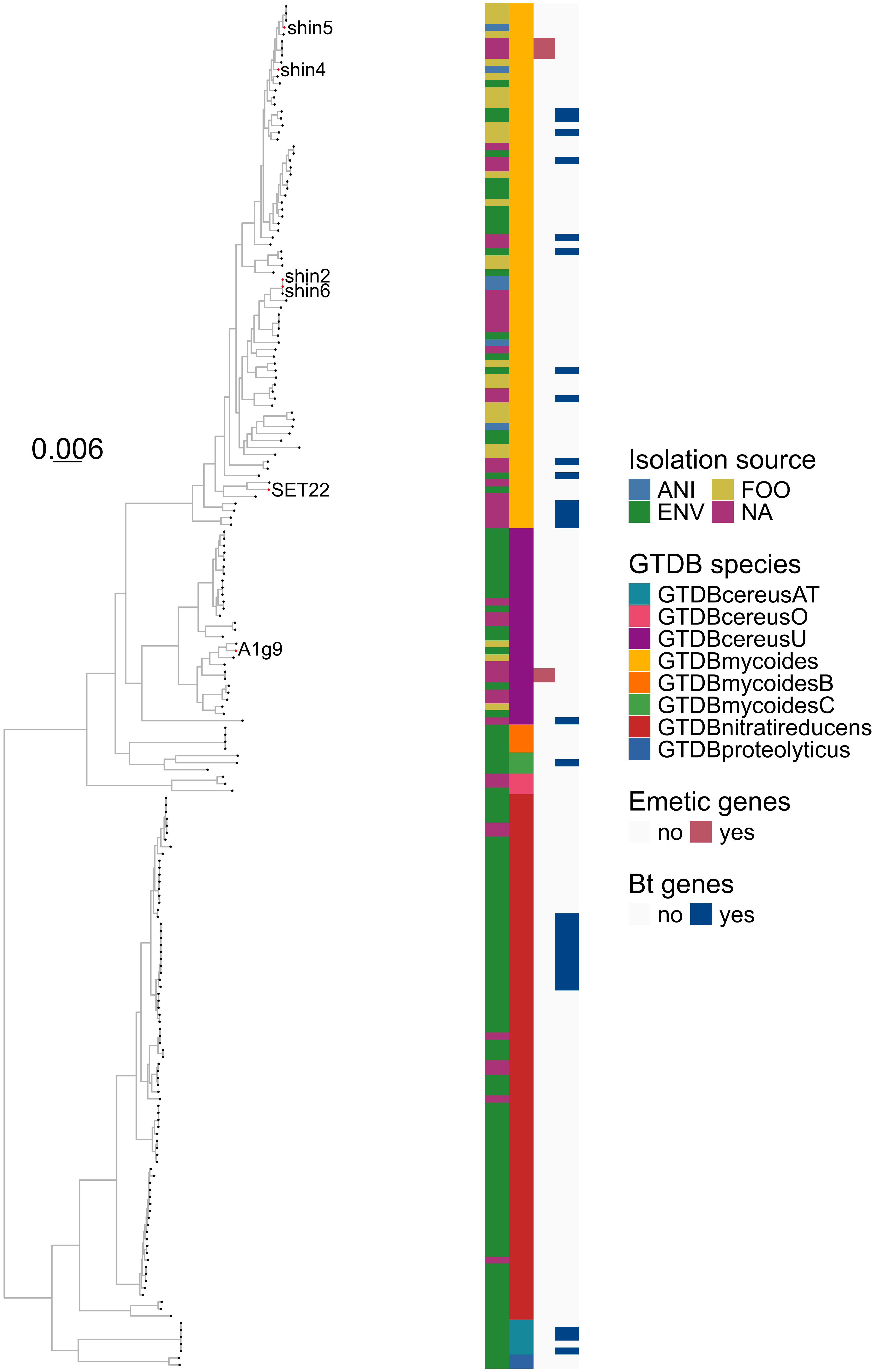

### Supplementary Figure S4

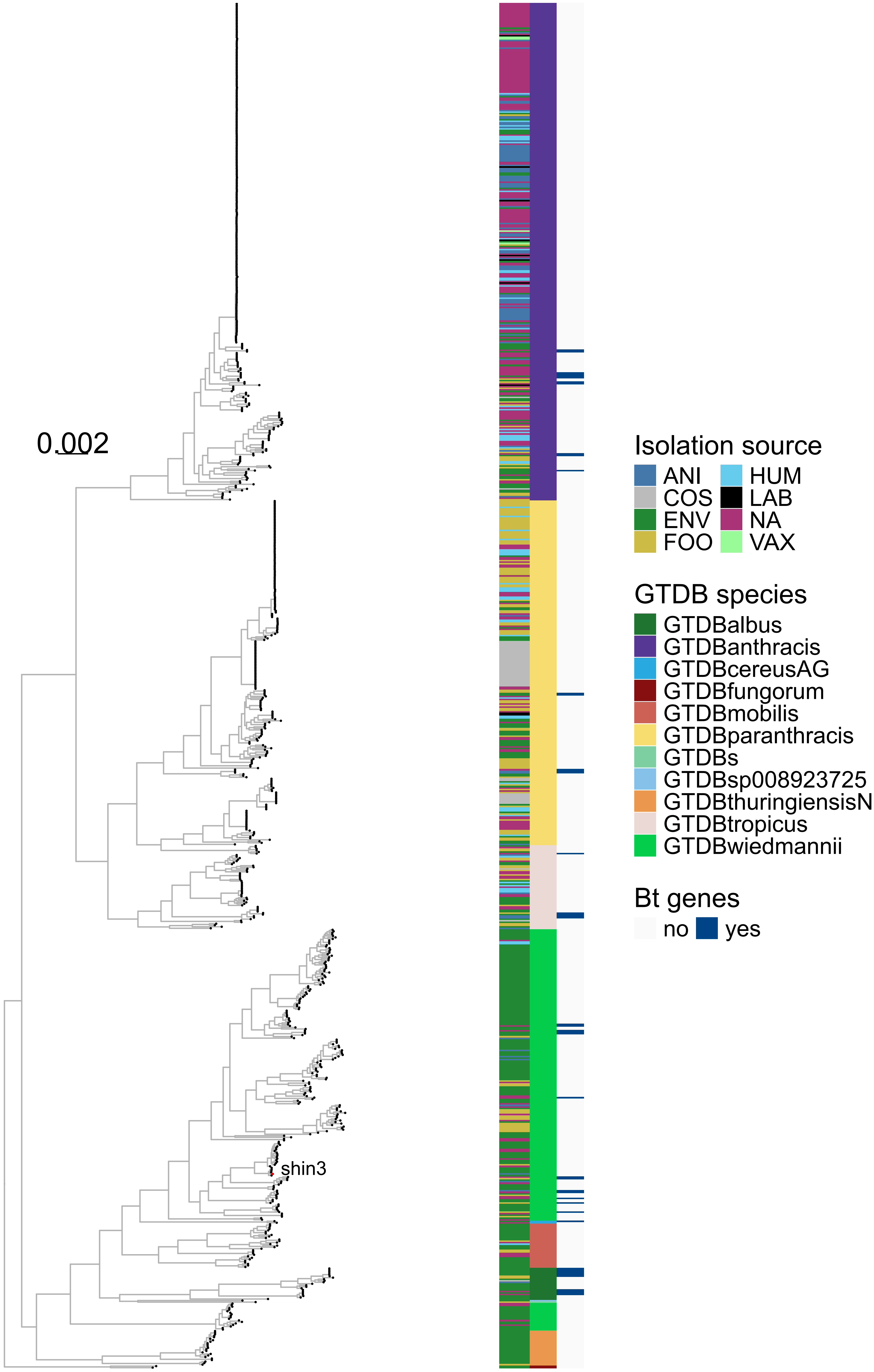

### Supplementary Figure S5

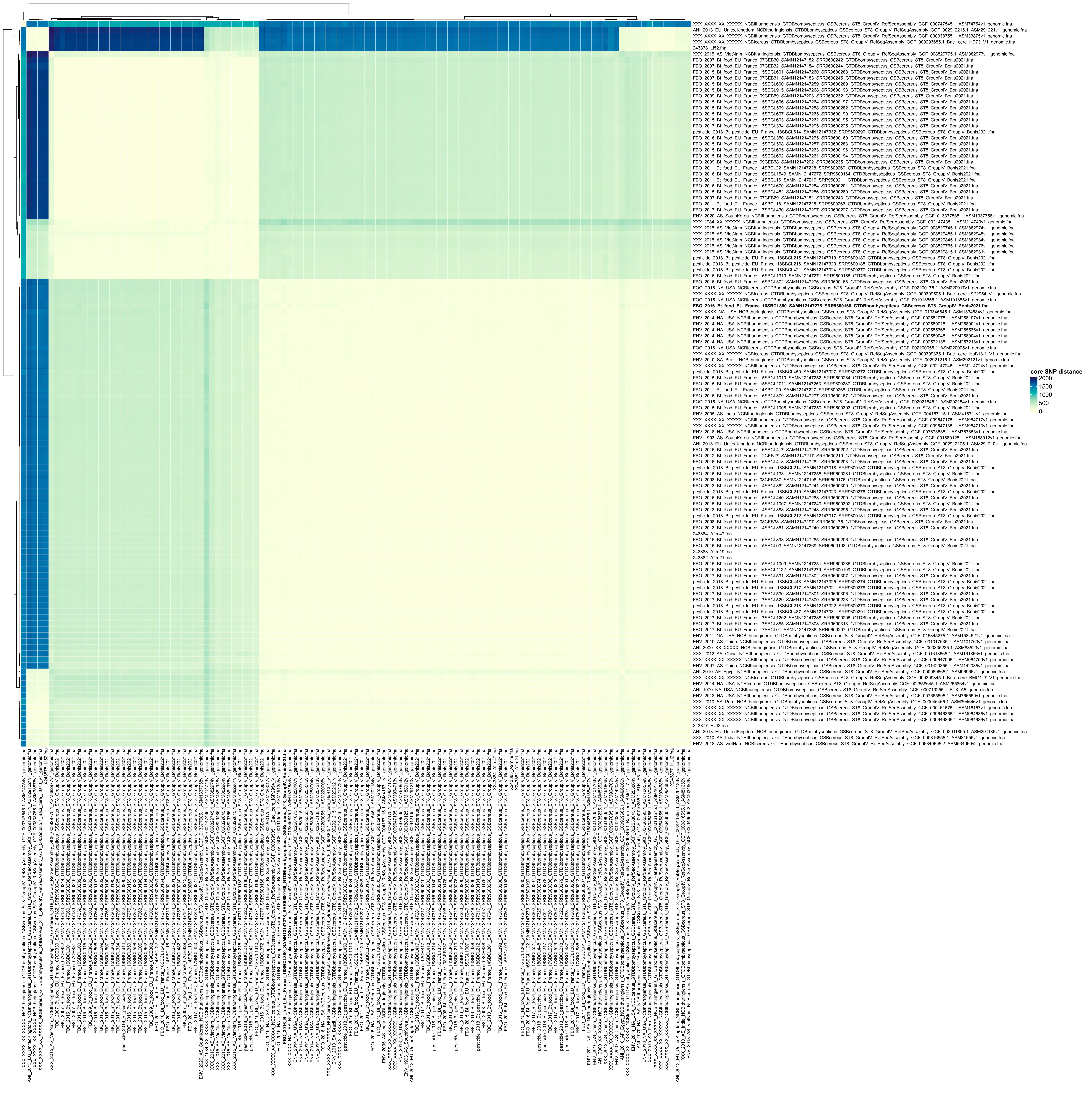

### Supplementary Figure S6

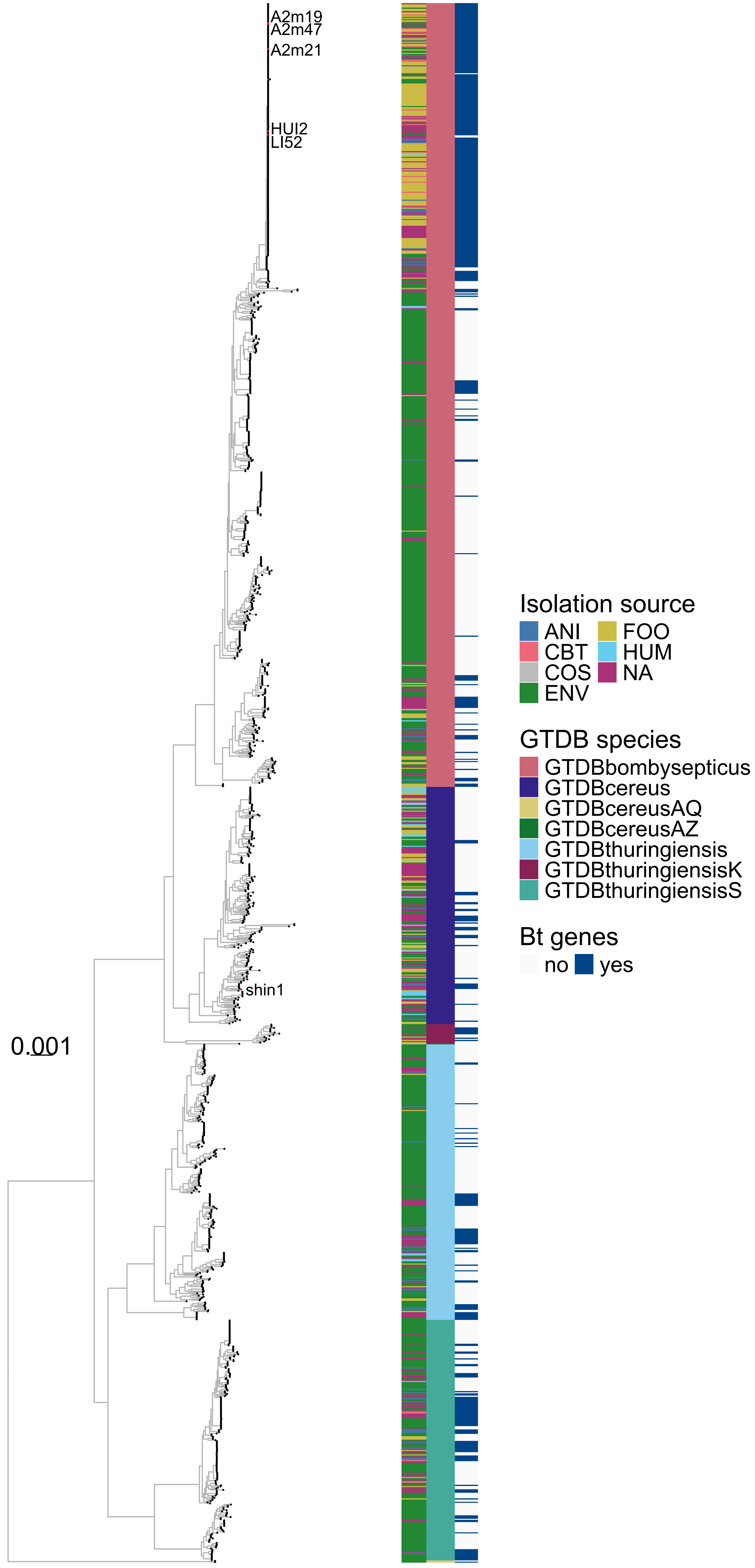

### Supplementary Figure S7

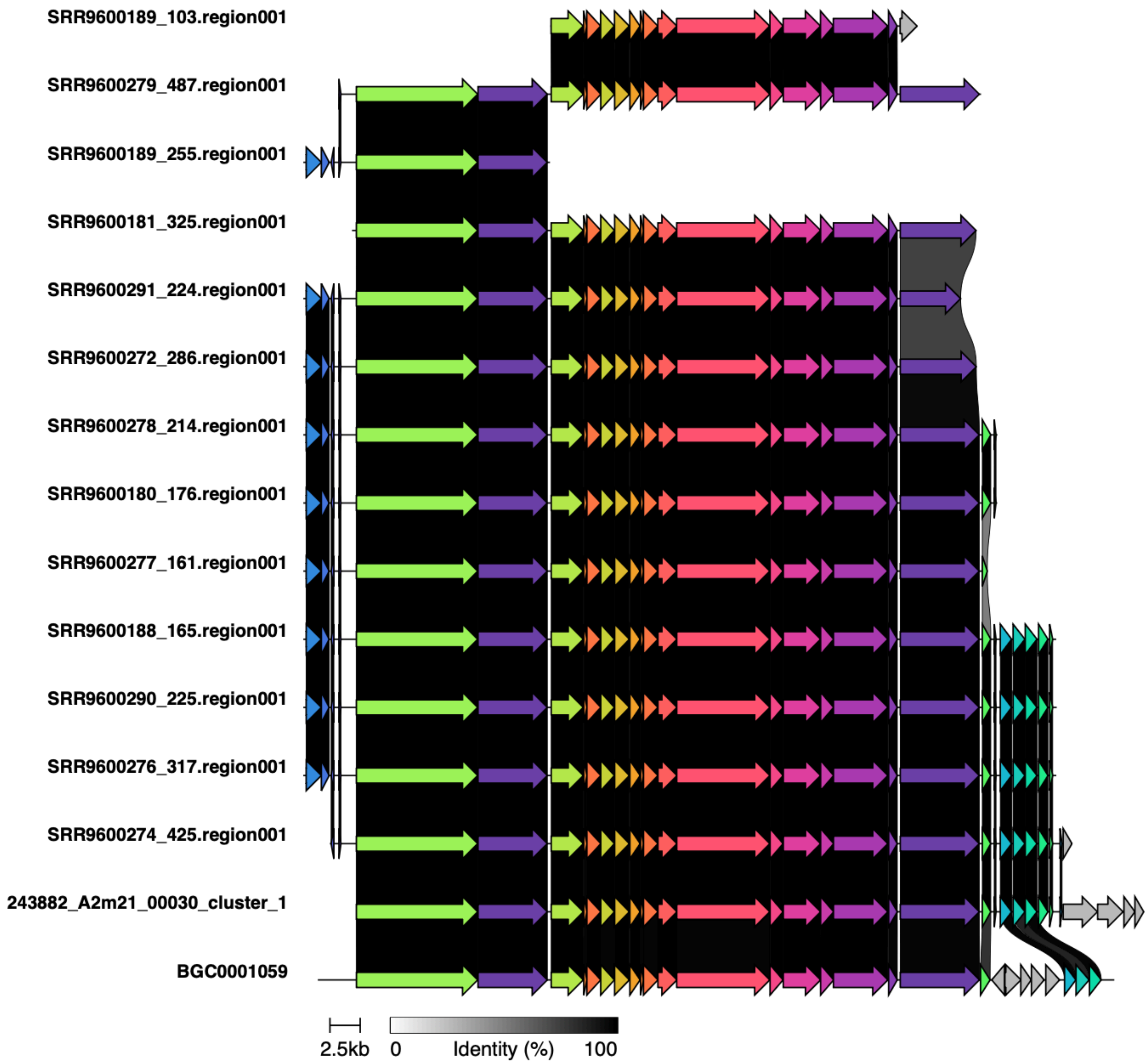
