## Supplementary figure legends for "THE INSECTICIDAL POTENTIAL OF *BACILLUS CEREUS* GROUP STRAINS FROM INSECT-DENSE REGIONS OF THE UNITED KINGDOM"

Supplementary Figure S1. Based on average nucleotide identity (ANI) distances, the novel strains are diverse. The branch lengths in the dendrogram represent ANI-distances, with numbers having been rounded to three significant figures. Only branch distances >0.01 are shown.

Supplementary Figure S2. An outlier is present among the ST8 genomes. Branch lengths represent dissimilarity (100-ANI (average nucleotide identity)), with the yellow vertical line representing an ANI distance of 99.5. The dendrogram was generated using bactaxR (v0.2.3; Jain et al. 2018; Carroll, Wiedmann and Kovac 2020).

Supplementary Figure S3. Maximum-likelihood phylogeny of *panC* Group VI and VIII. Colour strips to the right of the phylogeny denote (from left to right): (i) isolation source ("Isolation source"), (ii) Genome Taxonomy Database (GTDB) species ("GTDB species"), (iii) presence/absence of cereulide synthetase-encoding genes detected by BTyper3 ("Emetic genes"), (iv) presence/absence of *Bt*-associated toxin genes detected by BTyper3 ("Bt genes"). The phylogeny was rooted using the outgroup (*B. wiedmannii* RefSeq Assembly GCF_001583695.1). Branch lengths are reported in substitutions per site, and red tips denote novel strains investigated in this study. The numbers in the novel strain names have been omitted for legibility. "ANI" = animal-associated isolate; "FOO" = food isolate; "ENV" = environmental isolate; "NA" = not available; "COS" = cosmetics and consumer product isolate.

Supplementary Figure S4. Maximum-likelihood phylogeny of *panC* Group II and III. Colour strips to the right of the phylogeny denote (from left to right): (i) isolation source ("Isolation source"), (ii) Genome Taxonomy Database (GTDB) species ("GTDB species"), (iii) presence/absence of *Bt*-associated toxin genes detected by BTyper3 ("Bt genes"). The phylogeny was rooted using the outgroup (*B. cereus* RefSeq Assembly GCF_006094295.1). Branch lengths are reported in substitutions per site, and red tips denote novel strains investigated in this study. The numbers in the novel strain names have been omitted for legibility. "ANI" = animal-associated isolate; "FOO" = food isolate; "ENV" = environmental isolate; "NA" = not available; "COS" = cosmetics and consumer product isolate; "LAB" = laboratory strain; "VAX" = vaccine strain.

Supplementary Figure S5. SNP distance matrix of the ST8 genomes, including the novel strains. The cells are coloured based on the number of core SNPs, and the dendrograms represent the clustering of strains based on core SNPs using Euclidean distance.

Supplementary Figure S6. Maximum-likelihood phylogeny of *panC* Group IV. An expanded version of Figure 3A, with no collapsed clades. "COS" = cosmetics and consumer products.

Supplementary Figure S7. Clinker of zwittermicin A-like BGCs detected in commercial biopesticidal strains. Also included is the MIBiG zwittermicin A representative (BGC0001059) and a representative BGC from our novel strains (243882_A2m21_00030_cluster_1). Gene similarity (in % identity) is indicated by the shading.
